# NOS2 and COX2 Provide Key Spatial Targets that Determine Outcome in ER-Breast Cancer

**DOI:** 10.1101/2023.12.21.572859

**Authors:** Lisa A Ridnour, William F Heinz, Robert YS Cheng, Adelaide L Wink, Noemi Kedei, Milind Pore, Fatima Imtiaz, Elise L Femino, Ana L Gonzalez, Leandro Coutinho, Donna Butcher, Elijah F Edmondson, M. Cristina Rangel, Robert J Kinders, Stanley Lipkowitz, Stephen TC Wong, Stephen K Anderson, Danial W McVicar, Xiaoxian Li, Sharon A Glynn, Timothy R Billiar, Jenny C Chang, Stephen M Hewitt, Stefan Ambs, Stephen J Lockett, David A Wink

**Affiliations:** Cancer Innovation Laboratory, Center for Cancer Research, National Cancer Institute, National Institutes of Health, Frederick, MD; Optical Microscopy and Analysis Laboratory, Frederick National Laboratory for Cancer Research; Leidos Biomedical Research Inc. for the National Cancer Institute, Frederick, MD; Collaborative Protein Technology Resource (CPTR) Nanoscale Protein Analysis, OSTR, CCR, NCI, NIH; Imaging Mass Cytometry Frederick National Laboratory for Cancer Research.; Molecular Histopathology Laboratories, Leidos Biomedical Research Inc. for the National Cancer Institute.; Department of Pathology and Laboratory Medicine, Emory University, Atlanta, GA; Center for Translational Research in Oncology, ICESP/HC, Faculdade de Medicina da Universidade de São Paulo and Comprehensive Center for Precision Oncology, Universidade de São Paulo, São Paulo, SP, Brazil; Office of the Director, Division of Cancer Treatment and Diagnosis, NCI, Frederick, MD; Women’s Malignancies Branch, CCR, NCI, NIH; Houston Methodist Weill Cornell Medical College, Houston TX; Discipline of Pathology, Lambe Institute for Translational Research, School of Medicine, University of Galway, Galway, Ireland; Department of Surgery, University of Pittsburgh Medical Center, Pittsburgh, PA; Mary and Ron Neal Cancer Center, Houston Methodist Hospital and Weill Cornell Medicine, Houston, TX; Laboratory of Pathology CCR, NCI, NIH; Laboratory of Human Carcinogenesis, CCR, NCI, NIH, Bethesda, MD

**Author notes:** Co-senior author (Stephen J Lockett, David A Wink). Contributed equally (Lisa A Ridnour, William F Heinz, Robert YS Cheng, Adelaide L Wink).

## Abstract

Estrogen receptor-negative (ER-) breast cancer is an aggressive breast cancer subtype with limited therapeutic options. Upregulated expression of both inducible nitric oxide synthase (NOS2) and cyclo-oxygenase (COX2) in breast tumors predicts poor clinical outcomes. Signaling molecules released by these enzymes activate oncogenic pathways, driving cancer stemness, metastasis, and immune suppression. The influence of tumor NOS2/COX2 expression on the landscape of immune markers using multiplex fluorescence imaging of 21 ER-breast tumors were stratified for survival. A powerful relationship between tumor NOS2/COX2 expression and distinct CD8+ T cell phenotypes was observed at 5 years post-diagnosis. These results were confirmed in a validation cohort using gene expression data showing that ratios of NOS2 to CD8 and COX2 to CD8 are strongly associated with poor outcomes in high NOS2/COX2-expressing tumors. Importantly, multiplex imaging identified distinct CD8+ T cell phenotypes relative to tumor NOS2/COX2 expression in Deceased vs Alive patient tumors at 5-year survival. CD8+NOS2-COX2-phenotypes defined fully inflamed tumors with significantly elevated CD8+ T cell infiltration in Alive tumors expressing low NOS2/COX2. In contrast, two distinct phenotypes including inflamed CD8+NOS2+COX2+ regions with stroma-restricted CD8+ T cells and CD8-NOS2-COX2+ immune desert regions with abated CD8+ T cell penetration, were significantly elevated in Deceased tumors with high NOS2/COX2 expression. These results were supported by applying an unsupervised nonlinear dimensionality-reduction technique, UMAP, correlating specific spatial CD8/NOS2/COX2 expression patterns with patient survival. Moreover, spatial analysis of the CD44v6 and EpCAM cancer stem cell (CSC) markers within the CD8/NOS2/COX2 expression landscape revealed positive correlations between EpCAM and inflamed stroma-restricted CD8+NOS2+COX2+ phenotypes at the tumor/stroma interface in deceased patients. Also, positive correlations between CD44v6 and COX2 were identified in immune desert regions in deceased patients. Furthermore, migrating tumor cells were shown to occur only in the CD8-NOS2+COX2+ regions, identifying a metastatic hot spot. Taken together, this study shows the strength of spatial localization analyses of the CD8/NOS2/COX2 landscape, how it shapes the tumor immune microenvironment and the selection of aggressive tumor phenotypes in distinct regions that lead to poor clinical outcomes. This technique could be beneficial for describing tumor niches with increased aggressiveness that may respond to clinically available NOS2/COX2 inhibitors or immune-modulatory agents.

## Introduction

Breast cancer remains a leading malignancy in women. It is a heterogenous disease discerned by hormonal and HER2 status (1). While many treatment options are available for ER^+^ tumors, ER^-^ and triple negative breast cancers (TNBC) subtypes are the most clinically challenging malignancies due to limited treatment options (1). Recently, tumor NOS2 and COX2 co-expression has shown strong predictive power where their elevated expression correlates with reduced survival in ER-breast cancer (2). *In vitro* studies have identified several nitric oxide (NO)/PGE2 mechanisms that promote feed forward signaling loops leading to increased cancer stemness, metastasis, and immunosuppression (2–4). Importantly, immunosuppression is a key mechanism limiting cancer treatment efficacy (5). Many studies have shown that COX2-derived PGE2 and NOS2-derived NO promote immunosuppression in part by limiting activated T effector cells through increased Arg1, IL10, and TGFβ (6–9). In contrast, NOS2/COX2 blockade augmented cytolytic T cells, mature B cells, and neutrophils associated with resistance to 4T1 tumor rechallenge in 4T1 tumor bearing mice (10). Notably, these results are consistent with a recent phase 1/2 clinical trial demonstrating remodeling of the tumor immune microenvironment that involved increased B cells and neutrophils as well as improved overall survival and complete pathological responses in chemoresistant TNBC patients treated with taxol and the NOS inhibitor, L-NMMA (11).

Recent studies have shown that the spatial localization of CD8^+^ T cells is predictive of TNBC clinical outcomes where CD8^+^ T cell penetration into the tumor core (immune inflamed) correlated with pro-inflammatory immune response and predicted improved survival (12). In contrast, tumors with stroma-, margin-restricted, or abated CD8^+^ T cells (immune cold or immune desert, respectively) are immunosuppressed and predictive of poor clinical outcomes (12). As previously shown by us, elevated tumor COX2 expression may correlate with abated CD8^+^ T cell infiltration into breast tumor, whereas stroma restricted CD8^+^ T cells may correlate with both increased tumor NOS2 expression and tumor budding or satellitosis (13). Thus, we hypothesized that spatial relationships exist between tumor NOS2/COX2 expression and infiltrating CD8^+^ T cells that could influence clinical outcomes by promoting immunosuppression and increased disease aggressiveness.

Given that tumor NOS2/COX2 expression is antagonistic against CD8^+^ T cell function, herein we show significant relationships using the ratios of NOS2/CD8 and COX2/CD8 with respect to 5yr survival in ER-breast cancer patients (n= 21). Using spatial UMAP analysis we identify spatially distinct CD8^+/-^NOS2^+/-^COX2^+/-^ cellular neighborhoods with predictive value in these tumors including elevated fully inflamed CD8^+^NOS2^-^COX2^-^ cellular neighborhoods in surviving patients at 5yr. In contrast, CD8^-^NOS2^-^COX2^+^ immune desert regions were significantly elevated in deceased patient tumors. NOS2 was localized primarily at the tumor margins while COX2 expression extended deeper into the tumor core. Finally, the spatial analysis of CD44v6 and EpCAM cancer stem cell markers (SCS) within the context of tumor NOS2/COX2 expression identified cells that could contribute to stemness, drug resistance, and metastasis, which correlated with tumor NOS2/COX2 expression. Together, these observations describe candidate effects that spatially defined NOS2/COX2 expression patterns have in shaping the tumor immune microenvironment and the development of aggressive cancer phenotypes.

## Results

### Tumor NOS2/COX2 Expression and Survival

Previously, pathological IHC scoring revealed the power of elevated tumor NOS2/COX2 co-expression in predicting clinical outcomes in ER-breast cancer (2, 4, 14). Herein, we extend these earlier findings using multiplex immunofluorescence imaging, which facilitates the visualization and spatial localization of multiple targets at the single cell level. Spatial localization was determined using annotations of viable and necrotic tumor as well as tumor stroma regions that were annotated by a veterinary pathologist (EFE) on H&E images (QuPath) (15) and then fused with serial sections showing NOS2/COX2 fluorescent expression using HALO software. This methodology provides both quantitative and mechanistic descriptions of different cellular neighborhoods within the tumor microenvironment that correlate with disease progression. ER-breast tumors (n = 21 including 14 TNBC and 7 HER2-positive) from a previously reported cohort (2, 4, 14) were evaluated for the spatial localizations of tumor NOS2/COX2 expression in patients who survived (Alive) vs those who succumbed to disease (Deceased) at 5yr post-diagnosis. Consistent with the original pathologist IHC scores, the average NOS2/COX2 fluorescence cell intensities were significantly higher in Deceased vs Alive patient tumors (Fig. 1A-C) (4, 14). Quantitative analysis of NOS2/COX2 expressions revealed gradient changes in the signal intensity of NOS2 but not COX2. Therefore, NOS2/COX2 gradients were determined by adding 2, 4, or 6 standard deviations to the mean intensity to quantify weak, moderate, and strong tumor NOS2/COX2 signal intensities (Fig. 1C and Supplemental. Fig. 1A-C) (13). When stratifying for survival, the NOS2/COX2 signal intensities in tumors from Deceased vs Alive patients demonstrated a higher fold-change (∼6x) in the strong signal intensities of NOS2 (NOS2s) compared to moderate or weak (∼2-3x) signal intensities (Supplemental Fig. 1A). In contrast, the fold-change of the COX2 gradient was similar (2-3x) at all signal intensities (Supplemental Fig. 1B). These results suggest that NOS2 fluorescence signal intensity has higher predictive value than COX2, which is supported by NOS2 and COX2 HR of 6 and 2.5, respectively as reported by Glynn et al. (4, 14)

**Figure 1.**
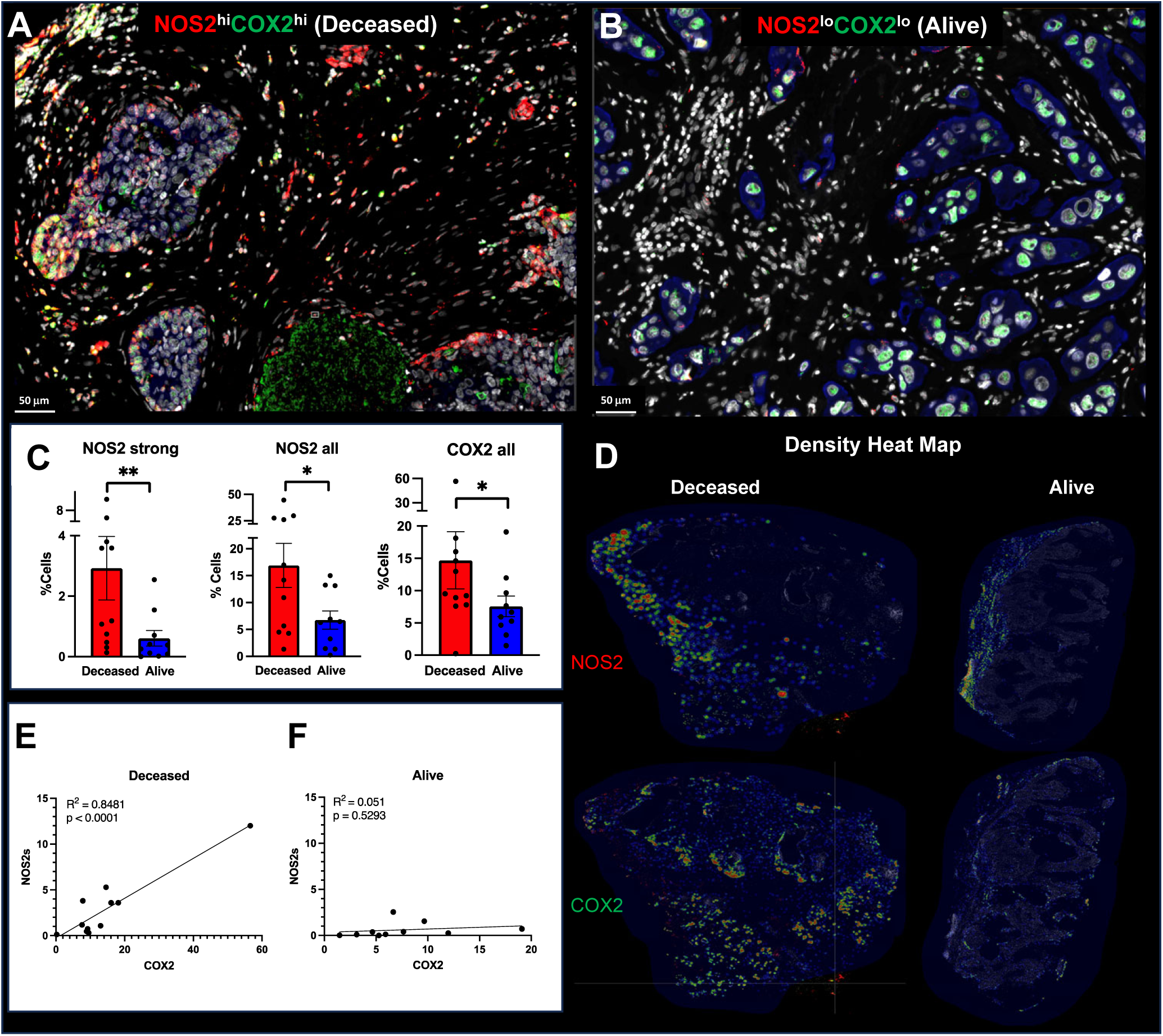
Spatial analysis of tumor NOS2/COX2 expression with respect to survival. Spatial landscape of NOS2 (red) and COX2 (green) with DAPI (white), the tumor marker CK-SOX10 (blue) for patient A) Deceased and B) Alive 5yr at survival. C) quantification of NOS2/COX2 tumor expression at the single cell level. D) heat density maps of tumor NOS2/ COX2 expression in Deceased and Alive patient tumors. E-F) Shows Pearson’s correlations between tumor NOS2 and COX2 expression R^2^ = 0.8481 and p<0.0001 in tumors from Deceased patients only.

Earlier studies identified NOS2/COX2 feedforward signaling that supports many oncogenic pathways in ER-breast cancer (2, 3). Moreover, a strong linear relationship between tumor NOS2/COX2 expression in all patient tumors has been reported (13). Given that the flux of NOS2-derived NO depends on the local density of NO producing cells as well as the concentration, diffusion, and reaction kinetics (16), we used density heat map analyses (Fig. 1D) to spatially quantify tumor NOS2/COX2 clustering in tumors from Deceased vs Alive patient tumors. Visual differences in density heat maps revealed increased spatially distinct tumor NOS2/COX2 clustering indicative of increased regional NO/PGE2 flux in Deceased patient tumors (Fig. 1D). Moreover, quantification of density heat maps showed significantly elevated NOS2 clustering in Deceased vs Alive patient tumors (Fig. 1D). A robust linear correlation was observed between strong NOS2 signal vs COX2 signal intensities in Deceased tumors (R^2^ = 0.8481, p < 0.0001) (Fig. 1E). In contrast, the NOS2/COX2 linear relationship in Alive tumors was not significant (R^2^ = 0.051, p = 0.5293) (Fig. 1F). These findings indicate that upregulated tumor NOS2/COX2 expression at the single cell level is predictive of poor survival. Thus, the single cell fluorescence analysis is consistent with the previous NOS2/COX2 IHC scoring, which showed strong predictive power of tumor NOS2/COX2 expression in ER-Breast cancer (2).

### Predictive Power of CD8^+^ T Cell Infiltration in the Context of Tumor NOS2/COX2

The spatial orientation of CD8^+^ T cells within the tumor landscape is predictive of TNBC survival (12). While the increased presence of CD8^+^ T cells in the tumor core is predictive of improved survival, their limited infiltration and/or stroma restriction correlated with poor clinical outcomes in TNBC (12). Given that 1) the NOS2/COX2 products NO/PGE2 are antagonistic to T effector cell function, 2) that tumor NOS2/COX2 expression limited CD8^+^ T cell penetration into the tumor core, and 3) that potent immune responses potentiate therapeutic efficacies, we reasoned that tumor NOS2/COX2 expression could influence therapeutic efficacies and clinical outcomes through their influence of CD8^+^ T cell penetration and function. Herein, we further explored this possibility by examining the spatial relationship between tumor NOS2/COX2 expression and CD8^+^ T cells in a breast cancer cohort at 5yr survival. The total numbers of CD8^+^ T cells and other immune markers did not change significantly between Deceased and Alive tumors in this cohort (Supplementary Fig. 2A). Our earlier findings demonstrated correlations between tumor NOS2/COX2 and the spatial orientation of CD8^+^ T cell in breast tumors, as well as tumor budding or satellitosis, which is an index of tumor invasiveness. We extended these earlier findings by examining the spatial localization of CD8^+^ T cells in Deceased vs Alive patient tumors (10, 13). Because NOS2/COX2 products NO/PGE2 have antagonistic effects on T effector cell function, NOS2/CD8 and COX2/CD8 ratios were quantified and found to be significantly elevated in tumors from Deceased patients, for both total and cytolytic (CD8^+^PD1^-^) CD8^+^ T cell populations (Fig. 2A). Moreover, PGE2 released from COX2 expressing tumor cells has been shown to impair the interaction of conventional dendritic cells (cDC1) with CD8^+^ T cells leading to decreased antigen presentation and recruitment of CD8^+^ T cells to the tumor microenvironment (17, 18). This possibility was explored in 4T1 tumor bearing mice treated with the clinically available NSAID indomethacin (INDO) that efficiently targets tumor COX2-expressing cells due to the slow rate of release of INDO from the COX2 enzyme (19, 20). In addition, INDO can also increase the PGE2 consumptive enzyme 15-PGDH (21). When compared to control untreated mice, INDO treatment led to increased expression of the cDC1 lineage determing factor IRF8, the c-type lectin-like activation receptor CLEC9a involved in anti-tumor immunity, chemokines CXCL9-11 that promote directional migration of immune cells, and IL27, which synergizes with IL12 to promote IFNγ production by CD4^+^, CD8^+^ T cells, and NKT cells (Supplemental Fig. 2A). The elevated NOS2/CD8 and COX2/CD8 relationships were validated in the Gene Expression Omnibus database inclusive for ER-breast cancer subtype, which showed significant hazard ratios (HR) of 5.67 and 3.34 at 5yr survival, for high vs low NOS2/CD8 and COX2/CD8 ratios dichotomized at the median, respectively (Fig. 2B). Importantly, these HR are consistent with those published earlier for tumor NOS2 (HR = 6.19) and COX2 (2.72) expressions (4, 14). These results suggest that elevated tumor NOS2/COX2 expression can suppress cytolytic CD8^+^ T effector cell populations during TNBC disease progression.

**Figure 2.**
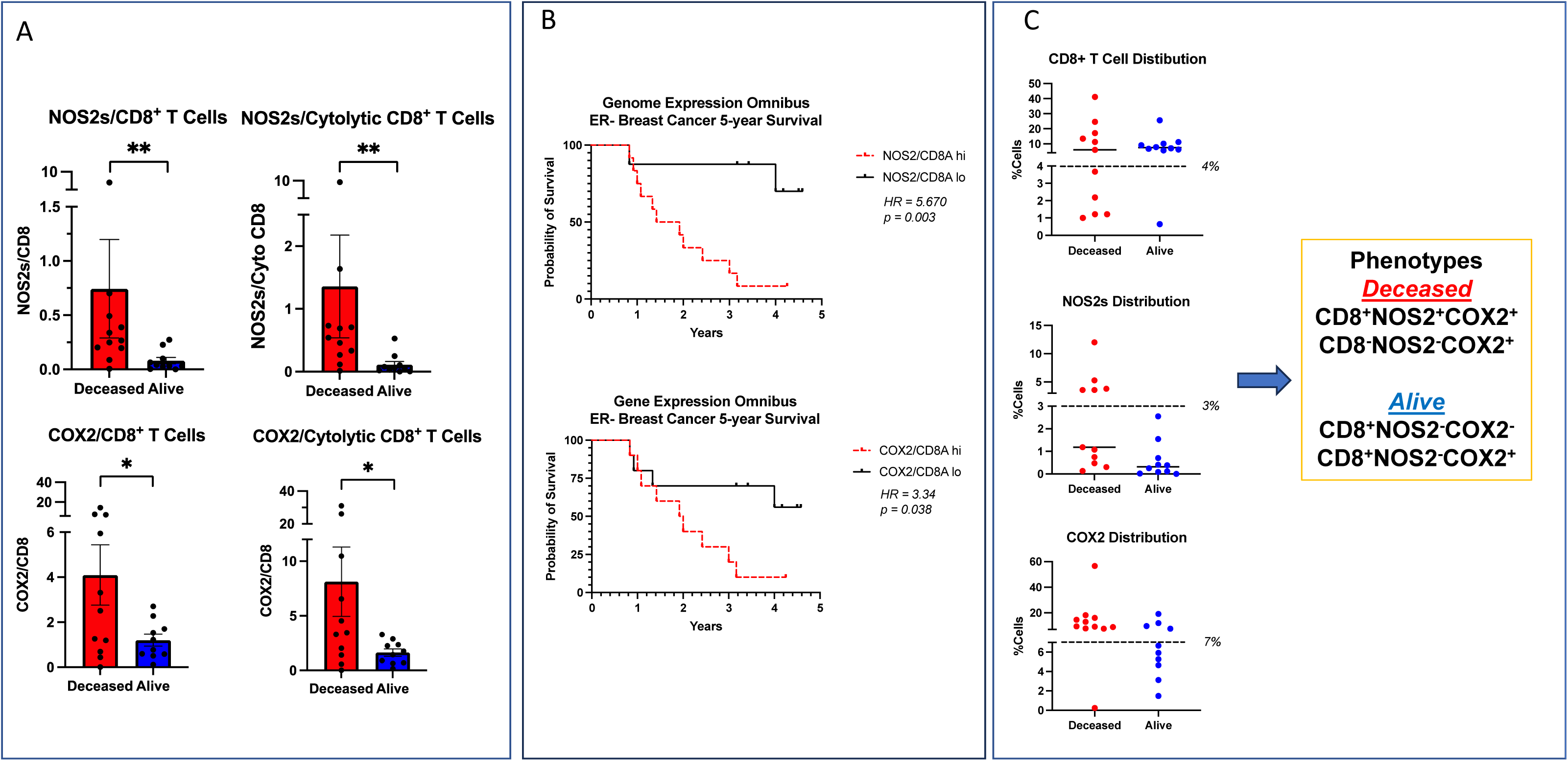
Analysis of tumor NOS2/COX2 expression with respect to CD8^+^ T cells. A) The ratio of NOS2s/CD8 and COX2/CD8 expression in total and CD3^+^CD8^+^PD1^-^ Teff cell populations. B) ER-breast cancer at 5yr survival stratified for NOS2/CD8 and COX2/CD8 ratios validation in GSE37751 breast cancer data from Genome Expression Omnibus (GEO) public data repository (https://www.ncbi.nlm.nih.gov/geo/info/download.html). The R software (version 4.2) was used to extract gene expression data from ER-samples for analysis of high (red) vs low (black) NOS2/CD8a and COX2/CD8a ratios dichotomized at the median. The survival data were exported to PRISM (version 9) and plotted for survival. Hazard ratios (HR) and p values were determined using Mantel-Haenszel and Gehan-Breslow-Wilcoxon test in PRISM software. C) Summarizes phenotype %Cell distributions in each tumor sample relative to survival. The dashed line denotes natural breaks in the data.

To further explore the impact of this relationship on clinical outcomes, tumor phenotypes were spatially defined relative to tumor NOS2/COX2 expression and the spatial localization of CD8^+^ T cells. The occurrence of these tumor phenotypes were compared between Deceased vs Alive patient tumors and summarized in Fig. 2C as %Cells for CD8^+^ T cell, NOS2s and COX2 distributions. Each distribution summary exhibits a natural break (black dashed line) separating high (+) vs low (-) marker expression; importantly, high T cell number or T cell abundance is definded as >4% (Fig. 2C). A comparison of the spatial localization of tumor phenotypes relative to NOS2/COX2 and CD8 expressions summarized in Fig. 2C and Supplemental Fig. 2B reveals two predominant phenotypes in Deceased vs Alive patient tumors, where CD8^+^NOS2^+^COX2^+^ (stroma restricted) and CD8^-^NOS2^-^COX2^+^ (immune desert) phenotypes were predominant in tumors from Deceased patients. In contrast, CD8^+^NOS2^-^COX2^-^ and CD8^+^NOS2^-^COX2^+^ fully inflamed phenotypes, where CD8^+^ T cells were observed to penetrate deep into the tumor cores and were present predominantly in Alive patient tumors (Fig. 2C and Supplemental Fig. 2B). The predominant phenotypes in the Deceased tumor samples were CD8^-^ with NOS2^-^ and COX2^+^ expression, which is defined as immune desert phenotype, while CD8^+^ with NOS2^+^ and COX2^+^ expression are defined as an inflamed stroma restricted phenotype, as determined by pathology, suggesting unique spatial configurations. Importantly, the common phenotype in Alive patient tumors was *CD8^+^NOS2^-^*, suggesting predictive power of this phenotype regarding clinical outcomes. To explore this possibility, NOS2s vs CD8 was visualized using a scatter plot (Supplemental Fig. 2C), which shows three distinct regions describing the influence of NOS2s on the CD8 status, including CD8^+^NOS2^-^ fully inflamed tumors with high CD8^+^ T cell tumor infiltration being present in 8 out of 10 tumors from Alive patients but only one Deceased patient tumor. In contrast, CD8^+^NOS2^+^ phenotypes were identified in inflamed regions where CD8^+^ T cells were spatially restricted in tumor stroma and high tumor NOS2 expression was observed at the tumor margin in 5 of 11 Deceased patient tumors. A CD8^-^NOS2^-^ immune desert phenotype commonly occurred in 5 out of 11 Deceased as well as 2 Alive patient tumors. Pearson’s correlation analysis (R^2^ = 0.54, p = 0.0241) of these results suggests a possible progression from CD8^-^NOS2^-^ immune deserts to CD8^+^NOS2^+^ restricted inflamed tumors (Supplemental Fig. 2D). These results show that tumors from surviving patients are highly inflamed with elevated infiltrating CD8^+^ T cells deep into the tumor core. In contrast, Deceased patient tumors are distinguished by two distinct phenotypes, cold immune desert regions and inflamed tumors with stroma restricted CD8^+^ T cells, where tumor NOS2 expression in the inflamed stroma determines outcome. Therefore, the spatial localization of NOS2 expressing tumor cells relative to CD8^+^ T cells could be key determinants of clinical outcomes.

### NOS2/COX2 Spatial Localization Defines the Tumor Immune Landscape

Previous studies have demonstrated the requirement of IFNγ and IL1/TNF cytokines for tumor NOS2/COX2 expression where correlations between tumor NOS2 and CD8 expressions were observed at the tumor stroma interface (3, 13, 22). To further explore this relationship relative to survival, spatial distribution and density heat maps were compared between Deceased vs Alive patient tumors, which demonstrated that tumor NOS2/COX2 and CD8^+^ T cell phenotypes occupy distinct regions in Deceased patient tumors (Fig. 3A). Comparison of NOS2 density heat maps in the deceased tumors show NOS2 expressing areas that are proximally orthogonal to COX2 and stroma restricted CD8^+^ T cells. In contrast, tumors from surviving patients exhibited reduced NOS2/COX2 clustering and increased CD8^+^ T cell penetration into the tumor core (Fig. 3 B). Thus, elevated tumor NOS2/COX2 expression in deceased patients is spatially distinct and associated with limited CD8^+^ T cell infiltration into the tumor core.

**Figure 3.**
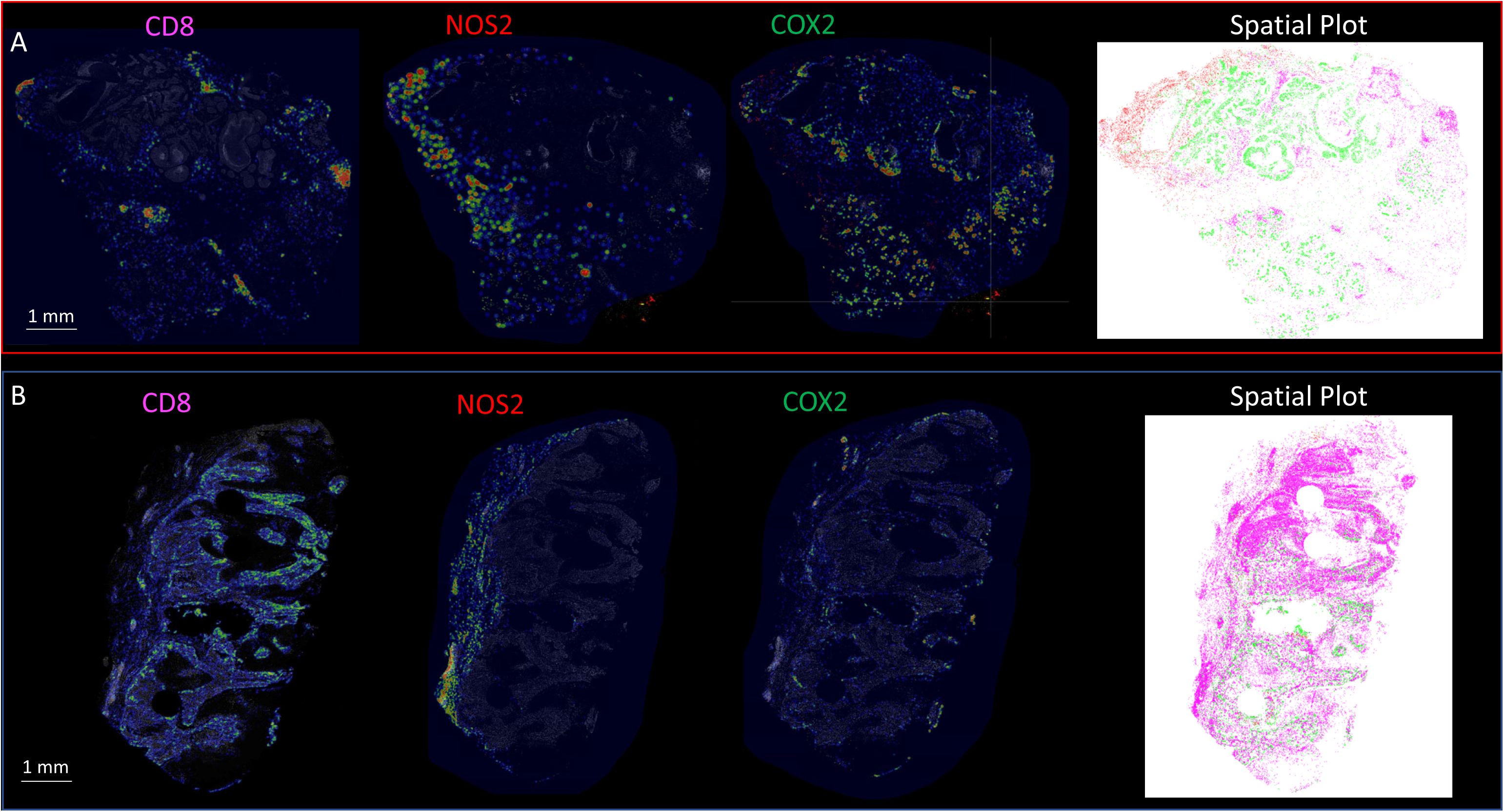
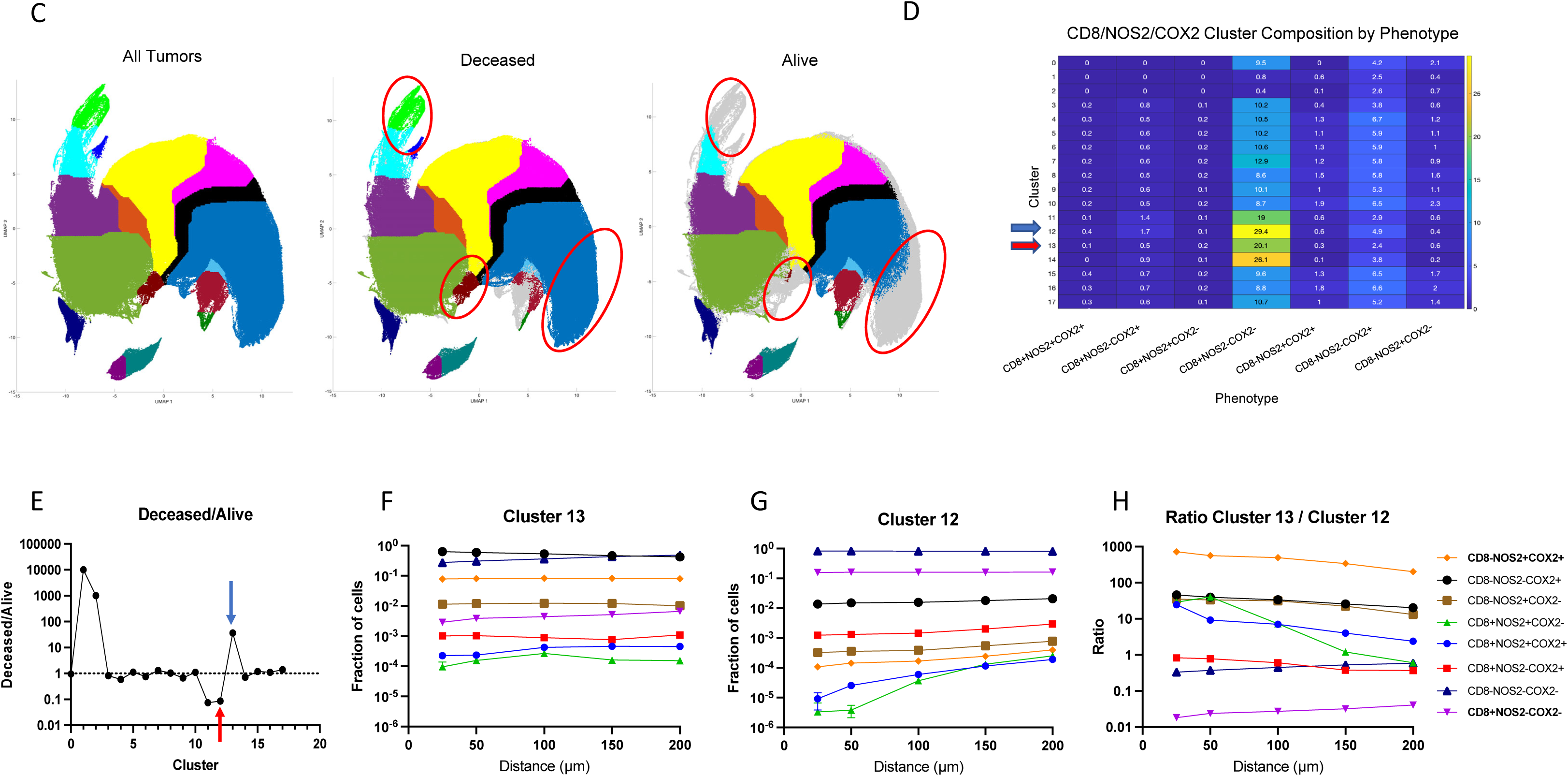
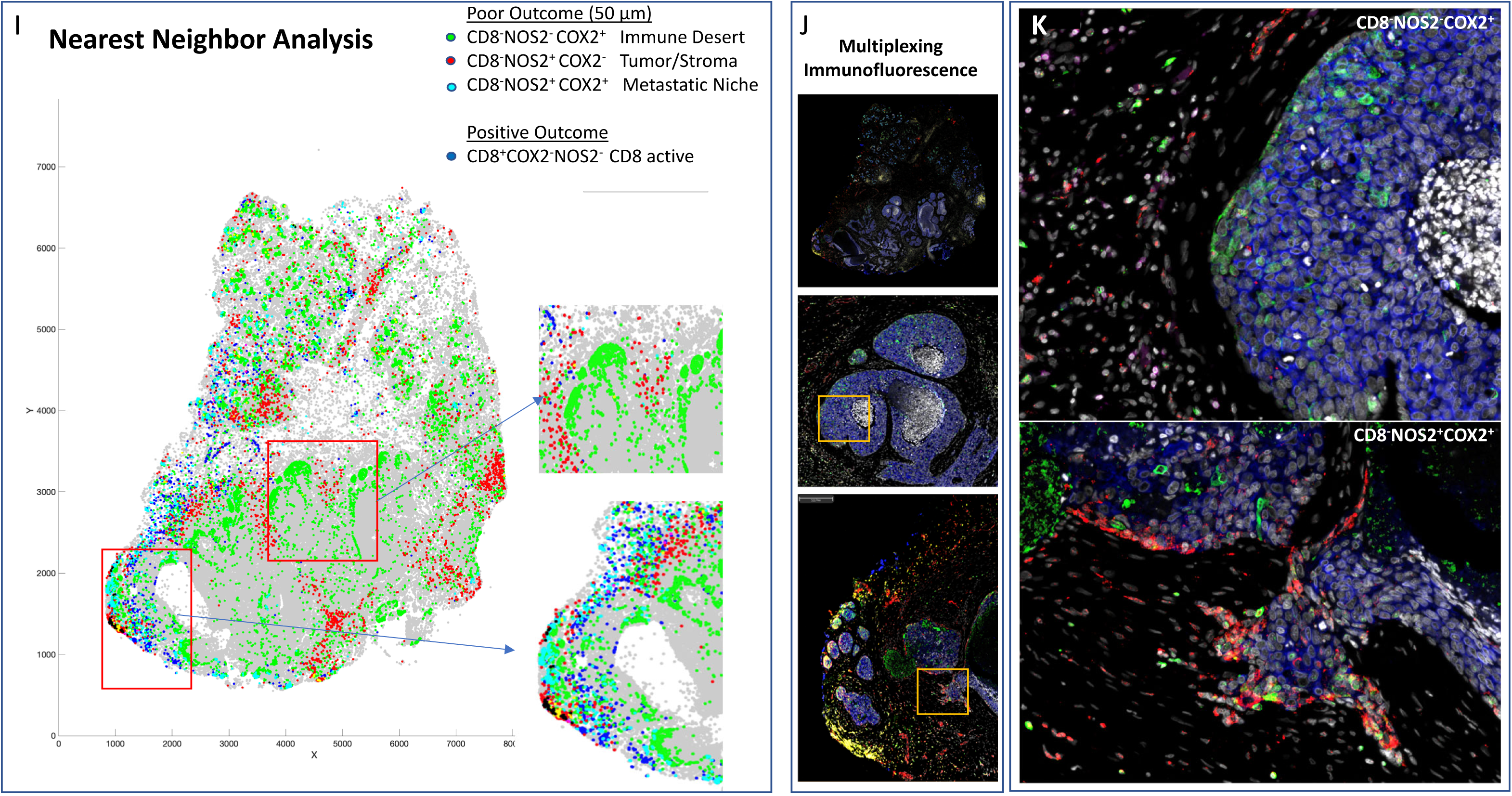
Spatial UMAP analysis for tumor NOS2/COX2 and CD8 expression. Heat density map comparison for NOS2, COX2 and CD8 expression in tumors from A) Deceased and B) Alive patients. Spatial distribution plots based upon positive pixels for NOS2 (red), COX2 (green) and CD8 (magenta) expression are shown. C) A spatial UMAP tumor analysis of cellular neighborhoods identifies differential neighborhood classes (clusters, red circles) in Deceased vs Alive patient tumors. D) Table summarizes the %cluster composition by each phenotype in all tumors. E) A comparison of cluster prevalence in Deceased vs Alice patients showing differential occurrence in clusters 12 and 13 in Deceased vs Alive patient tumors. F-G) Plots of the average Fraction of cells of each phenotype as a function of distance from each cell within the clusters that reveal distinct spatial distributions or density profiles of each phenotype. H) The ratio of phenotype density profiles for clusters 13 and 12 shown in panels F and G demonstrates the predictive value of CD8^-^NOS2^+^COX2^+^ and CD8^+^NOS2^-^COX2^-^ phenotypes due to their vast differences in Deceased vs Alive patient tumors. I) A spatial dot plot and nearest neighbor analysis at 50 μm distances of phenotypes with predictive value shown in panel H. J) Multiplex immunofluorescence of CD8 (magenta) NOS2 (red) and COX2 (green) expression in areas of interest (red square boxes) in the spatial dot plot shown in panel I. K) Enhanced magnification of areas of interest (yellow boxes) in panel J showing CD8^-^NOS2^-^COX2^+^ (immune desert, top) and metastatic niche associated with CD8^-^NOS2^+^COX2^+^ (inflamed region, bottom) phenotypes.

Next, these spatial configurations were quantified using spatial S-UMAP analysis. In contrast to the earlier visually guided analysis, the S-UMAP provides an unbiased, automated approach to define cellular configurations that predict survival. This approach was used to identify unique clusters that define cellular neighborhoods and nearest neighbors in the CD8^+/-^NOS2^+/-^ COX2^+/-^ phenotypes summarized in Fig. 2. Toward this end, the S-UMAP plots revealed distinct clusters of cellular neighborhoods in Deceased vs Alive patient tumors (Fig 3C), which was supported by cluster composition analysis that showed the density and distribution of CD8^+/-^ NOS2^+/-^COX2^+/-^ phenotypes at the whole tumor level (Fig. 3D). To identify cellular neighborhoods predictive of outcome, we plotted the ratio of the percentage of cells assigned to each cluster in Deceased vs Alive patient tumors. While most ratios were 1.0 and not predictive, three distinct clusters that increase in the tumors from Deceased patients (Fig. 3E blue arrows designating cluster #1, #2, and #13) and two clusters that increase in Alive patient tumors (Fig. 3E red arrows designating cluster #11 and #12) were identified. Only one of the three clusters (#13) that increased in Deceased patient tumors occurred in multiple samples, while the two clusters that increased in Alive tumors (#11 and #12) are very similar to each other. In tumors from Deceased patients the predominant phenotype was CD8^-^NOS2^-^COX2^+^, which corresponds to immune desert regions. In contrast, CD8^+^NOS2^-^COX2^-^ was the predominant phenotype in tumors from Alive patients, which corresponds with fully inflamed infiltrating CD8^+^ T cells that penetrated deep into the tumor core. These findings corroborate the phenotypes described above (Supplemental Fig. 2B-D) and support the idea that CD8^+^ T cell status impacts outcome, which is influenced by the spatial landscape of tumor NOS2/COX2 expression. Importantly, the unbiased S-UMAP approach comes to the same conclusions as the observer-driven approach and are thus supportive of one another.

The predictive value of clusters #13 and #12, which are cellular niches defined by the S-UMAP, was examined next. Plots of the average Fraction of cells of each phenotype as a function of distance from each cell within the cluster reveal distinct spatial distributions or density profiles of these phenotypes (Fig.3F and 3G). Importantly this analysis demonstrates the dominance of CD8^-^NOS2^+^COX2^+^ phenotype in cluster 13 (Fig. 3F yellow line), which represents a metastatic niche as shown in lower panel of Fig. 3I-K. In contrast, the dominance of the CD8^+^NOS2^-^COX2^-^ phenotype in cluster #12 (Fig. 3G purple line) represents high CD8^+^ T cell density in the tumor core of low NOS2/COX2 expressing tumors (Supplemental Fig. 3C). The ratio of phenotype prevalence between cluster #13 and #12 niches visualizes the vast difference between neighborhood compositions in the metastatic niche vs fully inflamed tumors with high CD8^+^ T cell density (Fig. 3H). Moreover, lack of CD8^+^ T cell penetration from the tumor/stroma interface into the tumor core supports observations of increased stroma restricted CD8^+^ T cells as well as immune desert regions lacking CD8^+^ T cells in Deceased patient tumors (Fig. 3F and Supplemental. Fig. 3A and 3B, respectively). In addition to the metastatic niche that predicted poor clinical outcomes, stroma restricted CD8^+^NOS2^+^COX2^+^ and CD8^+^NOS2^+^COX2^-^ phenotypes are also greater than 10. While CD8^+^NOS2^+^COX2^+^ and CD8^+^NOS2^+^COX2^-^ phenotypes comprise a smaller percentage of cells in cluster #13, their presence could have predictive value. Therefore, the CD8^+^ status of these phenotypes in cluster #13 could indicate an increased presence of metastatic or stem cell niches near stroma restricted CD8^+^ T cells. This more focused approach defines a spatial relationship between tumor NOS2/COX2 expression and CD8^+^ T cells where restricted CD8^+^ T cells that are excluded from the tumor are associated with elevated tumor NOS2 expression (Fig. 3F and H), while abated CD8^+^ T cell penetration into the tumor core corresponds with tumor COX2 expressing immune desert regions. Taken together, these analyses show that tumor NOS2/COX2 and CD8^+^ T cell spatial orientation defines distinct cellular neighborhoods with predictive power.

To further explore the spatial significance of CD8^+^ T cells relative to tumor NOS2/COX2 were examined with respect to known pathological features of the tumor microenvironment. Fig. 4A-C shows five distinct annotated areas including 1) lymphoid aggregates (Fig. 4A orange circle); 2) areas of tumor fragmentation (< 0.1 mm^2^) (Fig. 4B); 3) regions associated with larger tumor nests (> 0.05 mm^2^) (Fig. 4A magenta circle); 4) the tumor core (Fig. 4A blue circle); 5) NOS2^+^ and NOS2^-^ tumor edges (Fig. 4C yellow box), defined as the region at the tumor interface of larger tumor nests. The lymphoid aggregates were conglomerates of CD3^+^ lymphoid cells at the tumor margin in tumors from both deceased and surviving patients. The CD8^+^ T cell distributions in these defined regions were examined in all tumors, where distribution analysis showed elevated CD8^+^ T cell populations in lymphoid aggregates while the lowest CD8^+^ T cell distribution was observed in the tumor core of immune desert regions (Fig. 4D). Further stratification for survival revealed that the CD8^+^ T cell distribution in Alive patient tumors was approximately 4-fold higher in the tumor core when compared to Deceased Patient tumors (Table I). Immune deserts were previously defined as 100 cells/mm^2^, which is approximately 1% cells (12). This again confirms that CD8^+^ T cells in Deceased patient tumors are restricted and marginalized. In contrast, increased CD8^+^ T cell infiltration was observed in Alive patient tumors. Thus, the result that CD8^+^ T cells are stroma restricted in Deceased patient tumors but highly infiltrate tumor cores in Alive patient tumors is consistent with the nearest neighbor analysis in the S-UMAP shown in Fig. 3 and Supplemental. Fig. 3.

**Figure 4.**
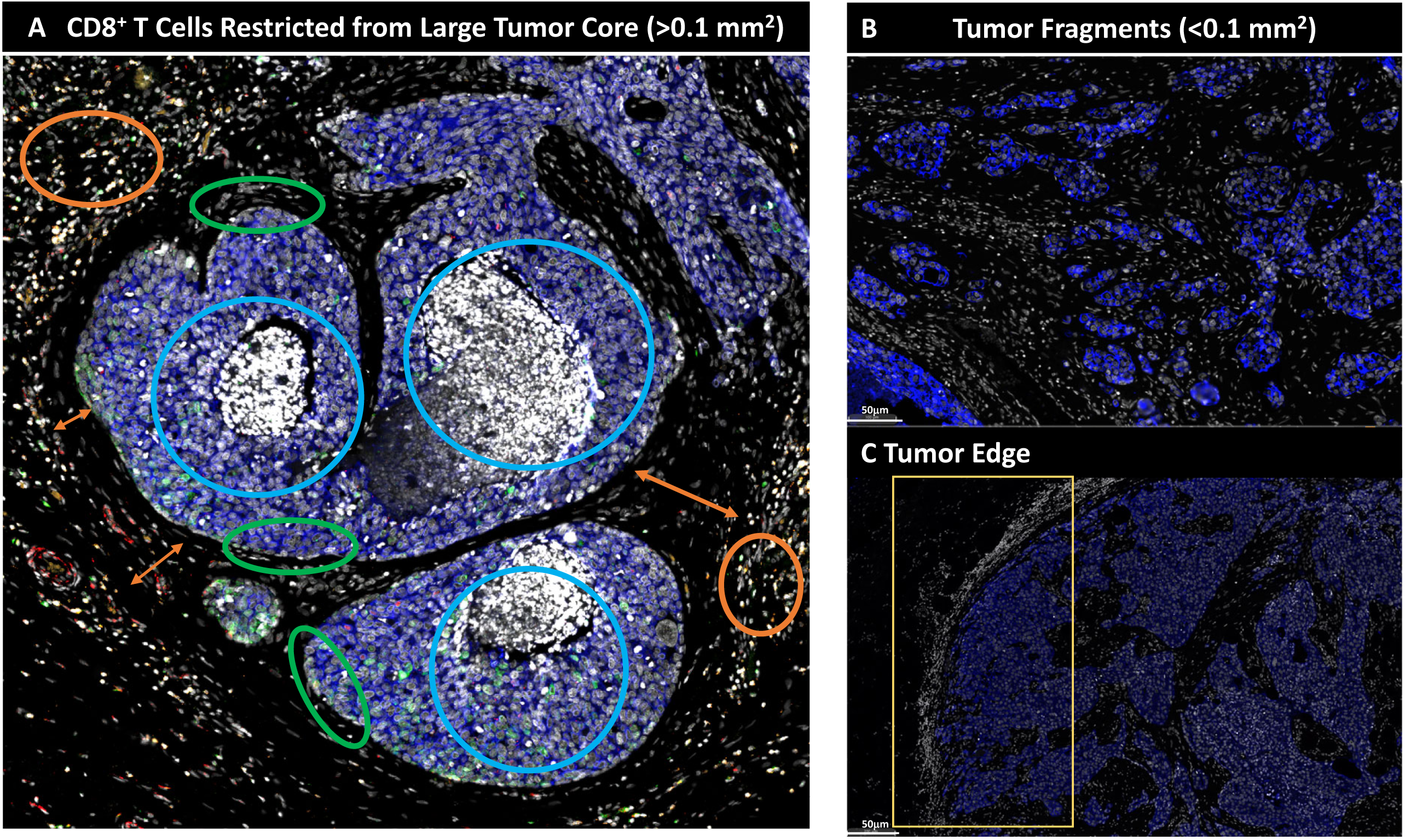
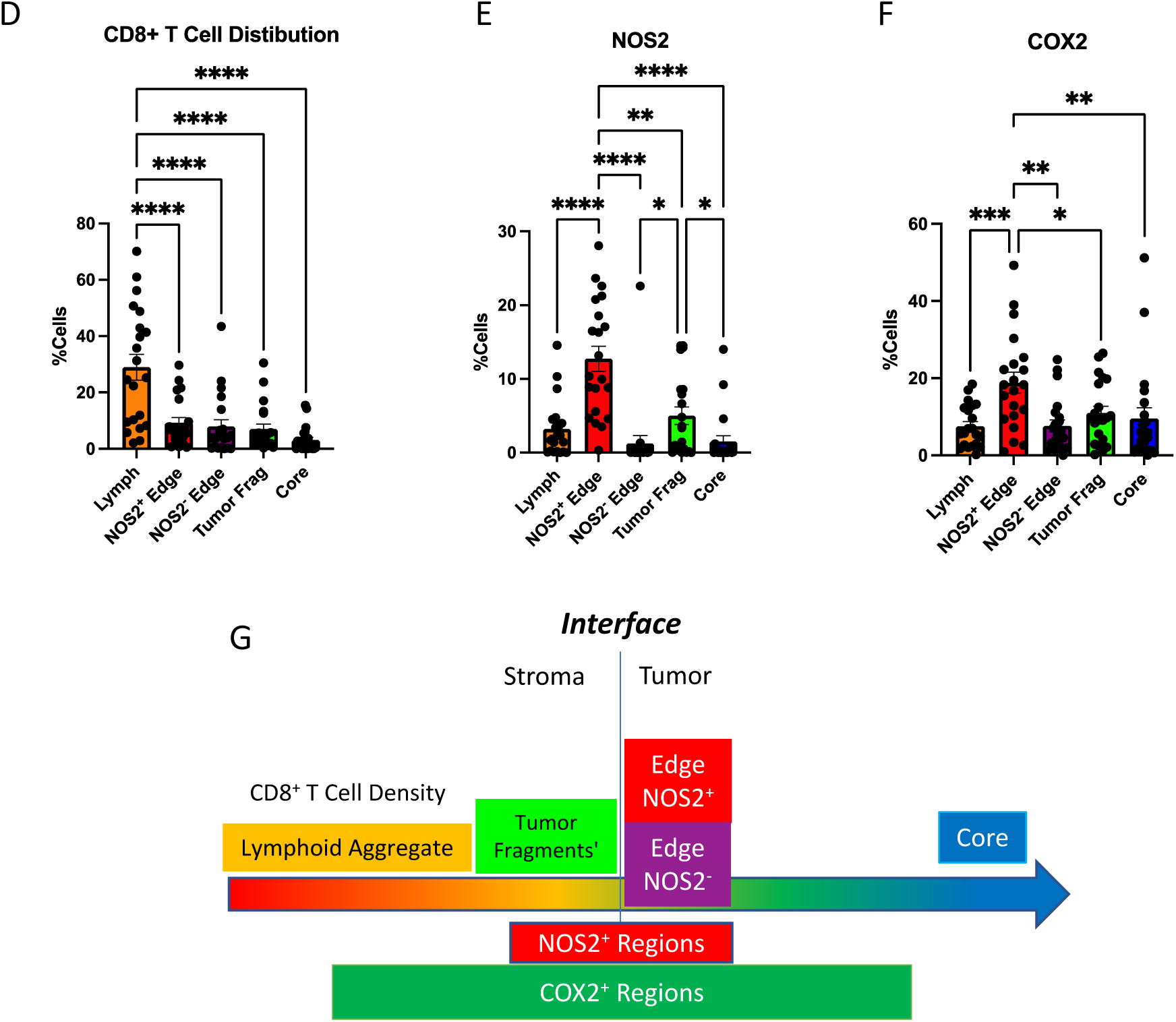
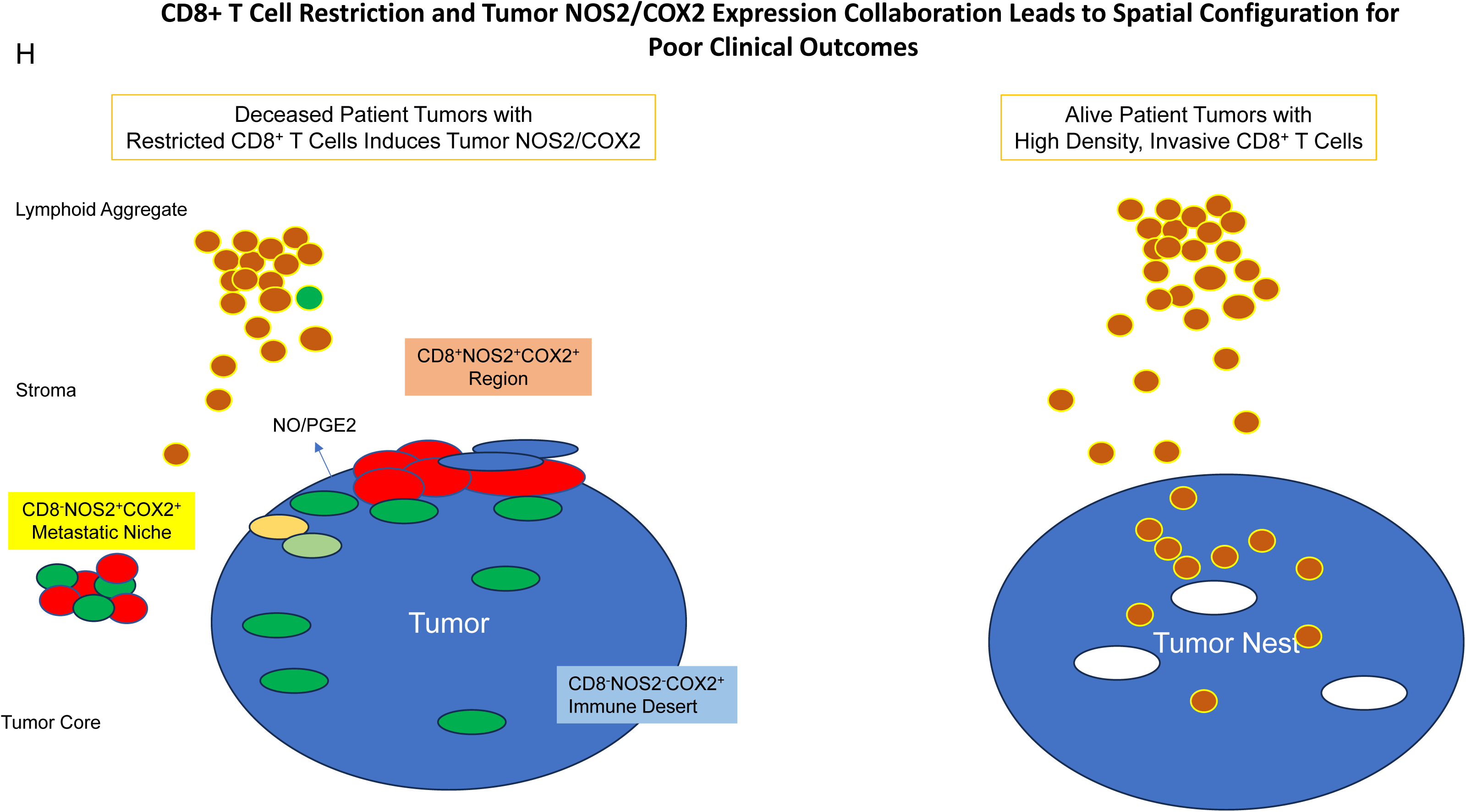
Defined CD8 NOS2 and COX2 spatial landscape. Five basic regions showing A) margin or stroma restricted lymphoid aggregates (orange circles) where a gap of 50 µm (double orange arrow) between CD3^+^ T cells aggregates and the tumor edge (green circle) was observed. Blue circles identify immune desert regions lacking CD8^+^ T cells. B) Tumor fragmentation or satellite region. C) Tumor edge with proximal stroma regions. Significant differences in %cell composition are shown for D) CD8^+^ T cells as well as E) NOS2^+^ and F) COX2^+^ tumor cells. G) Graphic summary of tumor NOS2/COX2 landscape and CD8^+^ T cell regional distributions with respect to Tumor-Stroma interface. H) Spatial architecture of predictive phenotypes in Deceased vs Alive patient tumors.

**Table I.** %CD3CD8^+^ T Cells in Designated Regions.

| Region | Deceased | Alive |
| --- | --- | --- |
| Lymph | 25 | 37 |
| Tumor Fragments | 5 | 10 |
| NOS2+ | 8.8 | 12 |
| NOS2- | 7 | 8 |
| Core | 1.0 | 4.4 |

The spatial localization of tumor NOS2/COX2 expression vs CD8^+^ T cells shows a different pattern within the defined regions. Tumor NOS2 density was evaluated at the tumor edge (Fig. 4E) where tumor satellite regions indicative of invasive tumor cells (13) contained significantly higher NOS2 expression while NOS2 was lowest in lymphoid aggregates and the tumor core (Fig. 4E). Importantly, the tumor core is nearly devoid of NOS2 (Fig. 4E) as well as CD8^+^ T cells, supporting the idea that NOS2 resides at the tumor/stroma interface as previously described (13). NOS2 was significantly higher in the tumor satellite regions when compared to the tumor core (Fig. 4E). In contrast, COX2 expression was more evenly distributed in the defined regions (Fig. 4F). COX2 was significantly higher in association with the NOS2^+^ tumor edge but was also found in the tumor core and stroma restricted lymphoid aggregates (Fig. 4F). The analysis shows clear distinction and regional distribution of CD8^+^ T cells and tumor NOS2/COX2 that correlates with survival (Fig. 4G, Fig. 1).

Tumors from Deceased patients exhibit gaps separating larger tumor nests from stroma restricted CD8^+^ T cells, which was not observed in tumors from surviving patients. As shown above, both NOS2 and COX2 can be expressed on larger tumors, and NOS2^+^/COX2^+^ foci are associated with areas of inflammation proximal to stroma restricted CD8^+^ T cells as previously described (13). Larger tumor nests (> 0.05 mm^2^) exhibit gaps (average of 50 µm) between tumor and lymphoid cells (Fig 4A orange arrows). Stroma restricted lymphoid aggregates average 500-1000 μm from NOS2^+^ and/or COX2^+^ tumor edges. Stroma restricted CD8^+^ T cells were observed 50-100 µm from the tumor margin of NOS2^-^COX2^+^ tumor edges. While COX2 is expressed at the edge of immune deserts, its expression is abated deeper at the core of immune deserts. These observations could indicate that COX2/PGE2 may serve as a barrier preventing CD8^+^ T cell infiltration into the tumor core, thus facilitating the development of an immune desert, which is consistent with increased CD8^+^ T cell penetration to the core associated with COX inhibition by NSAID treatment (10). These observations further support a role of tumor NOS2/COX2 expression during progression from inflamed foci to immune desert regions.

The above results show two principle CD8^+^ T cell spatial orientations associated with tumor NOS2/COX2 expression, which are inflamed, restricted lymphoid restricted and immune deserts devoid of lymphoid cells (Fig. 4H). The immune desert is COX2^+^ but NOS2^-^ however, NOS2^+^ and COX2^+^ tumor satellites form in the inflamed areas near stroma-restricted CD8^+^ T cells that produce IFNγ (13). Importantly, these NOS2^+^ tumor satellites exhibit increased elongation and migration consistent with increased tumor metastatic potential (13). Elevated tumor NOS2/COX2 promote a feed forward mechanism that drives cancer cell phenotypes with metastatic, chemoresistant, and cancer stem cell (CSC) properties (3, 23). Tumor CD44v6 and EpCAM expression have been used as clinical CSC markers in breast and other cancers, correlating with metastasis, circulating tumor cells, CSC, and chemoresistance (24). Next, these markers were used to identify CSC niches in the tumor microenvironment (TME) during disease progression.

Spatial localization of tumor CD44v6 and EpCAM expression showed distinct expression patterns (Fig. 5A). Univariant analysis demonstrated significantly higher CD44v6 expression in tumors from surviving patients while EpCAM did not change significantly with respect to survival (Fig. 5B CD44v6 all, EpCAM all). Given that CD44v6 is expressed on lymphocytes and modulates their functional activity (25) and that tumor NOS2/COX2 expression modulates CD8^+^ T cell penetration into the tumor core (10), potential roles for tumor NOS2/COX2 relative to CD44v6 and EpCAM expression were evaluated by comparing ratios of tumor NOS2/COX2 to CD44v6 and EpCAM. Toward this end, the ratios of NOS2/CD44v6, NOS2/EpCAM, COX2/CD44v6, and COX2/EpCAM were significantly elevated in Deceased patient tumors when compared to Alive patient tumors as shown in Fig. 5B. These results suggest that elevated tumor NOS2/COX2 could influence the effects of EpCAM and CD44v6 relative to clinical outcomes. We recently demonstrated significant correlations between tumor NOS2/COX2 expression and both stroma restricted CD8^+^T effector cells and secreted IFNγ (13). Next, we explored potential associations between stroma restricted CD8^+^T effector cells and IFNγ identified in lymphoid aggregates with the cancer stemness EpCAM, and CD44v6 using Pearson’s correlation coefficients, where significant R^2^ values were determined for T effector cells or IFNγ vs EpCAM in tumors from Deceased patients (Fig. 5D). In contrast, significant correlations were not observed for CD44v6 as this biomarker was largely identified is immune desert regions devoid of CD8^+^ T cells. The significant correlations identified between CD8^+^T effector cells, IFNγ and EpCAM suggests that EpCAM could be induced by INFγ in lymphoid aggregate regions as summarized in Fig. 5E. To explore this possibility, the spatial geography of EpCAM as well as CD44v6 was examined using information from the S-UMAP analysis shown in Fig. 3G, which identified regional CD8^-^ NOS2^+^COX2^+^, CD8^-^NOS2^+^COX2^-^ and CD8^-^NOS2^+^COX2^+^ cellular phenotypes in Deceased patient tumors. As shown in Fig. 6A, EpCAM is expressed on the tumor edge in NOS2^+^COX2^+^ niches (Fig. 3G), while CD44v6 is predominantly localized in COX2^+^ immune desert regions of the tumor core (Fig. 3G). The S-UMAP analysis revealed highly significant correlations between EpCAM in lymphoid restricted regions where elevated IFNγ and NOS2 were expressed, either at the tumor edge or in tumor satellites areas (Fig. 6B). While EpCAM is expressed with sporadic CD44v6 expression in lymphoid aggregate areas, EpCAM expression is absent along tumor NOS2^-^ edges. These results suggest a progression from inflammatory NOS2^+^ regions where EpCAM is predominant to immune desert regions expressing COX2 and CD44v6.

**Figure 5.**
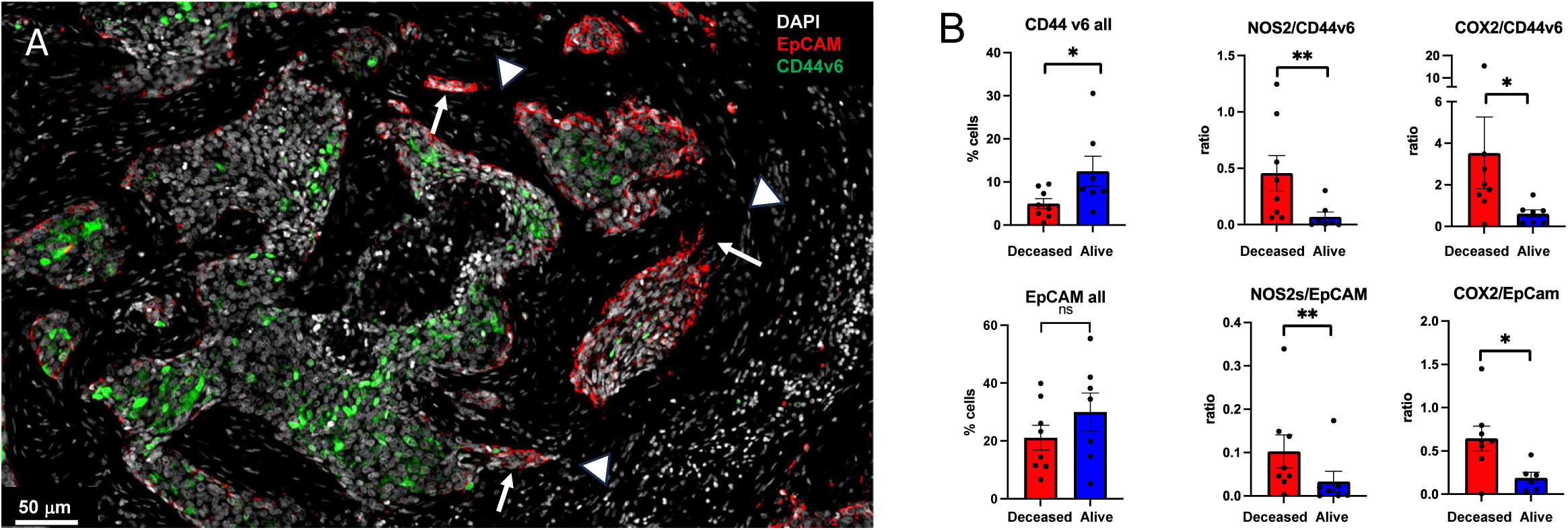
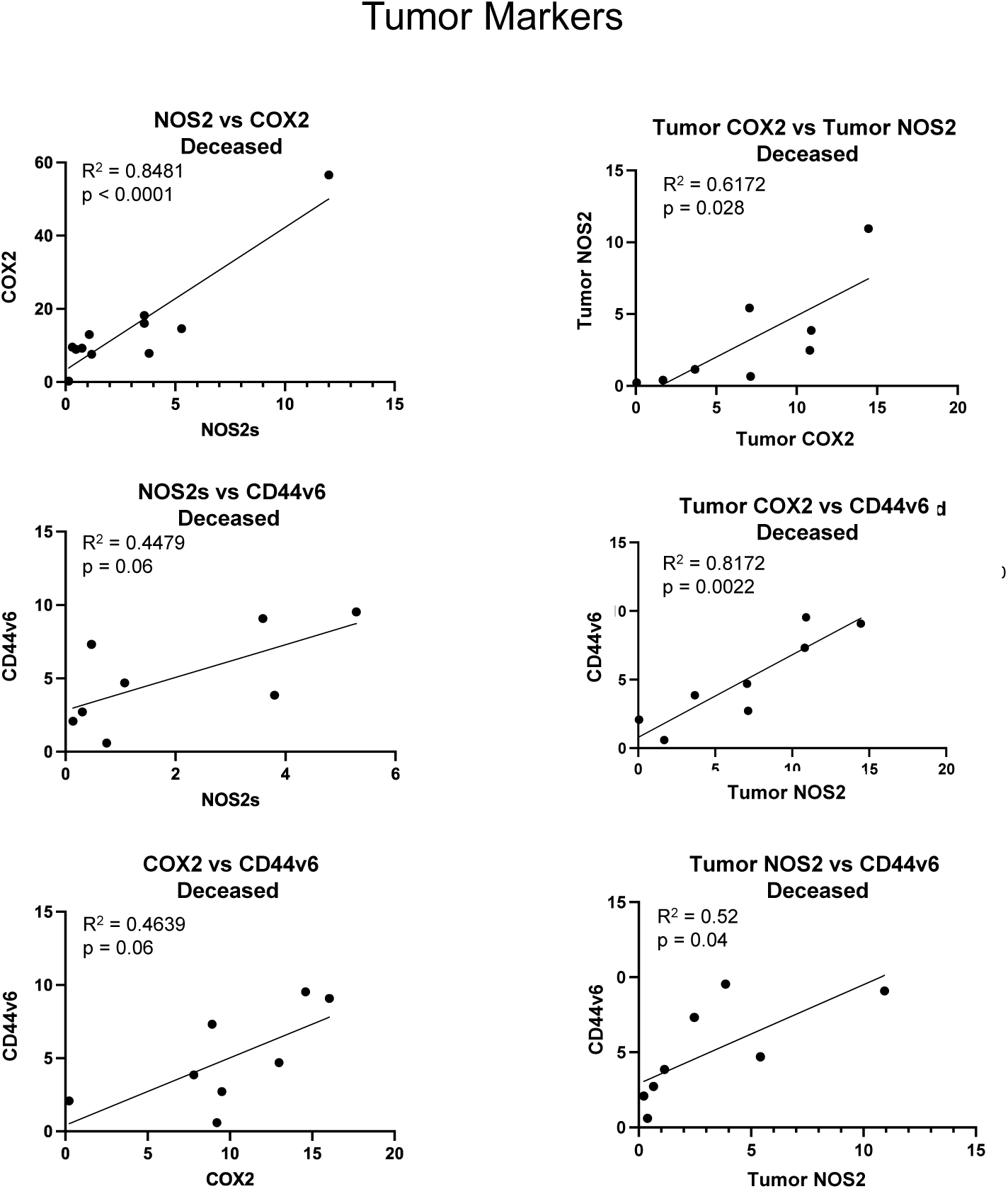
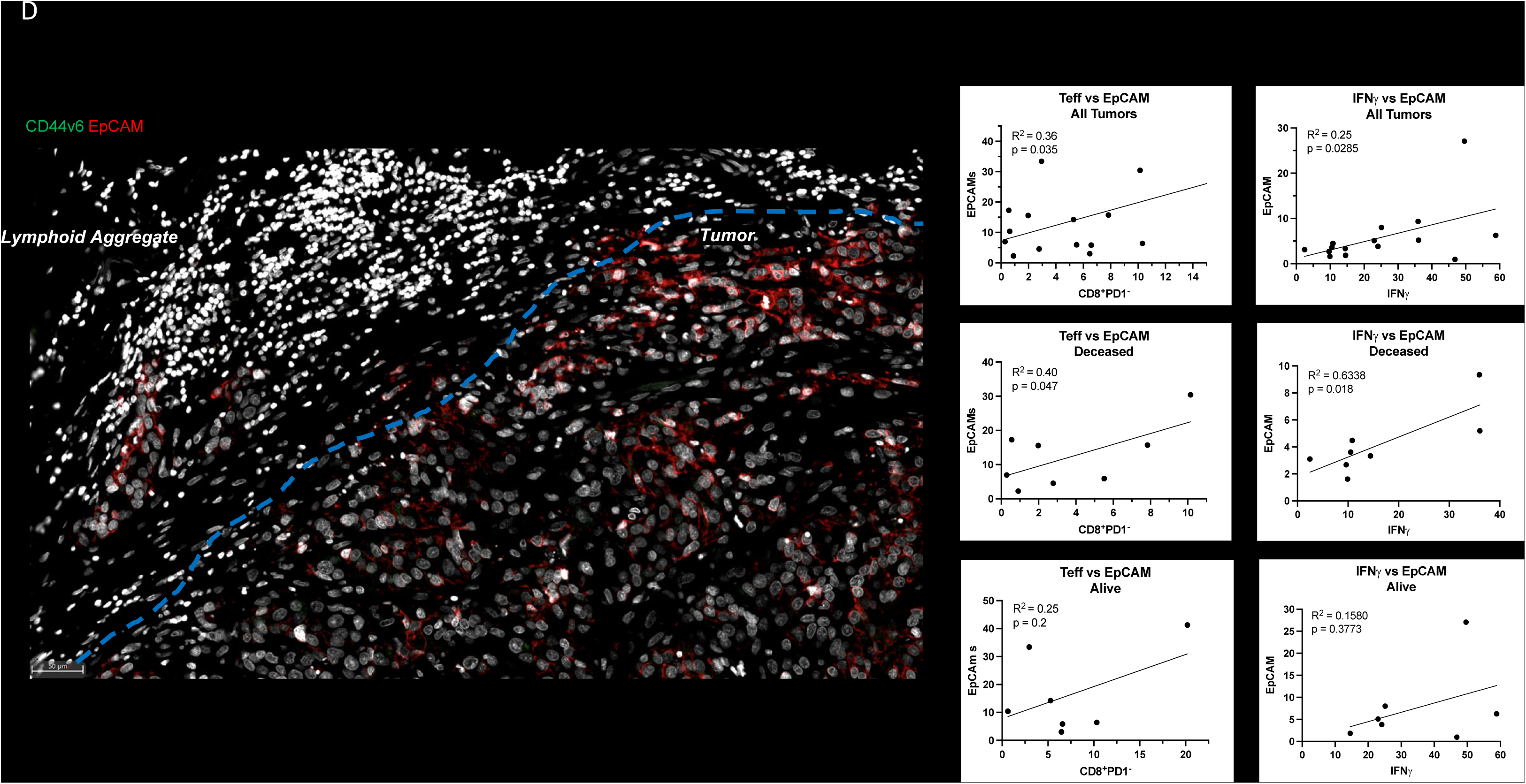
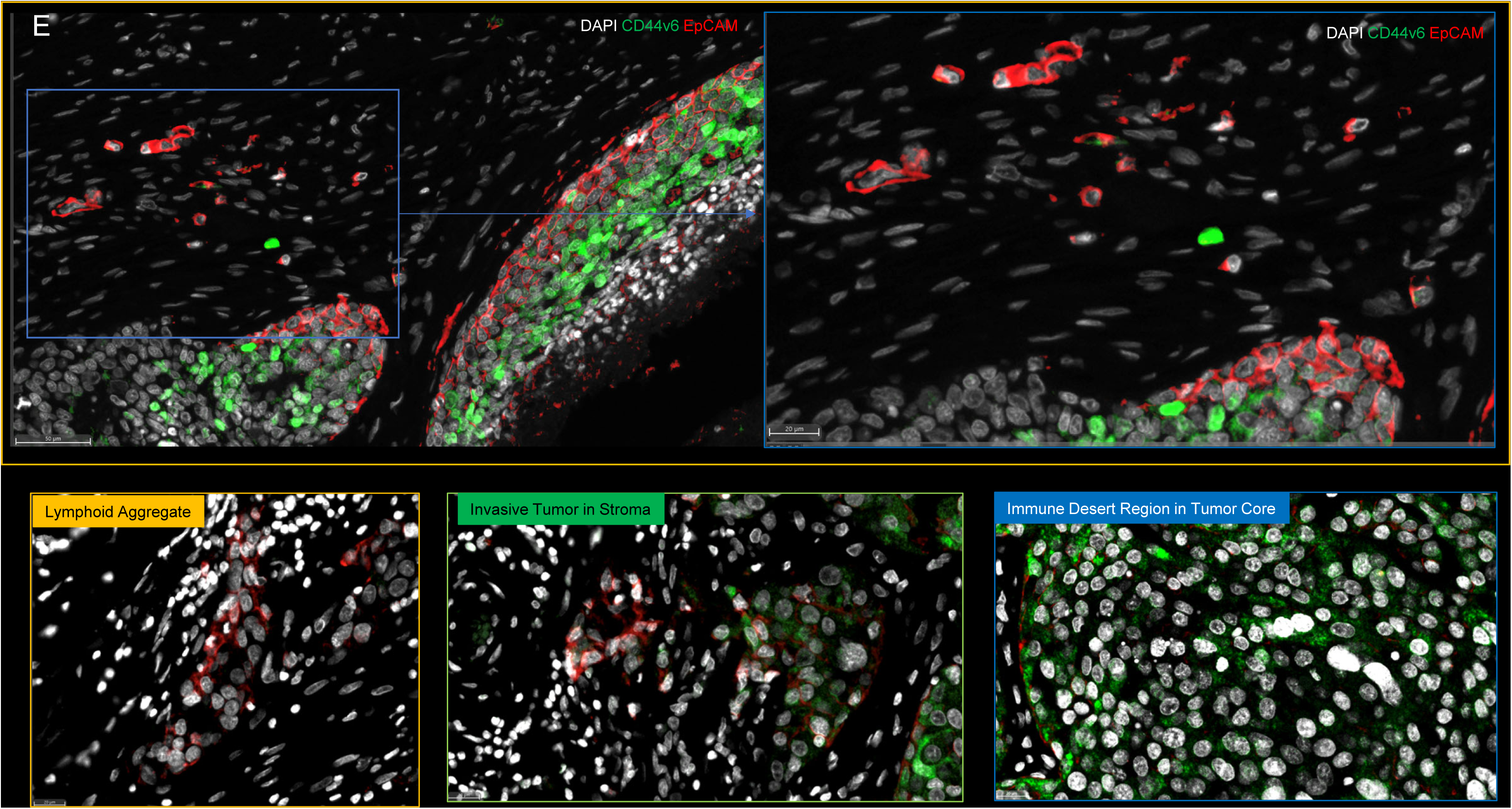
Spatial landscape of CD44v6 and EpCAM expression relative to tumor NOS2/COX2 expression and survival. A) Enriched regions showing spatially distinct CD44v6 and EpCAM expression. B) Quantification of CD44v6 and EpCAM as well as ratios of NOS2/COX2 to CD44v6/EpCAM, respectively, in Deceased vs Alive tumors. Significance was determined using Mann Whitney test where * P < 0.05 and ** P = 0.007. C) Pearson’s correlation coefficient showing significant associations between NOS2/COX2 and CD44v6 expression in Deceased vs. Alive patient tumors. D) Pearson’s correlation coefficient showing significant associations between T effector cell or IFNγ and EpCAM expression in Deceased vs. Alive patient tumors. E) Spatial landscape of CD44v6 and EpCAM expression in different annotated regions.

**Figure 6.**
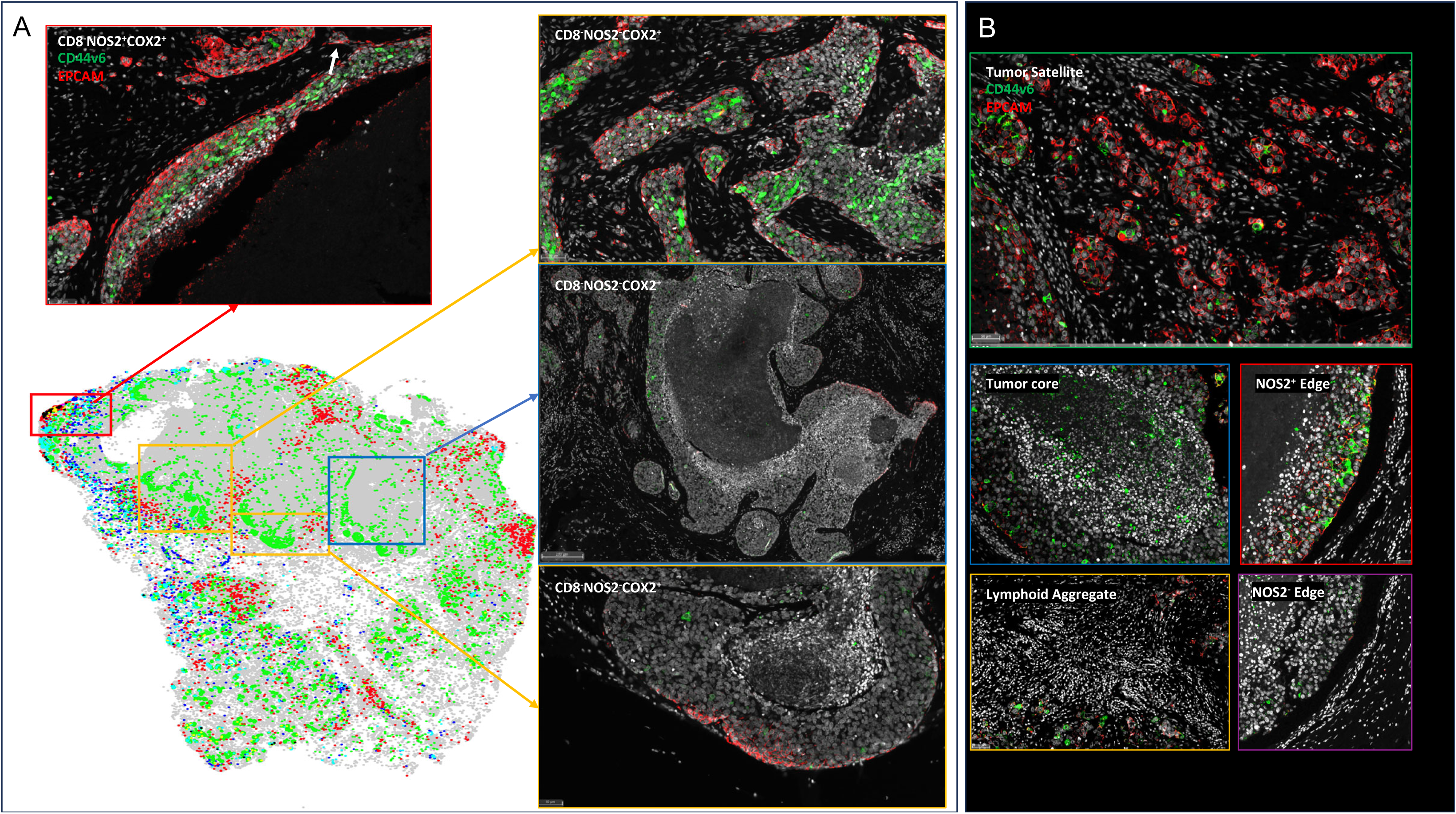
S-UMAP and regional annotations to identify cellular neighborhoods of interest with respect to the tumor cancer stem cell markers EpCAM and CD44v6. A) Unsupervised analysis of S-UMAP highlighting specific magnified regions; red box showing CD8^-^NOS2^+^COX2^+^ phenotype in stroma restricted inflamed region; orange and blue boxes showing CD8^-^NOS2^-^COX2^+^ phenotypes in immune desert regions. B) Supervised analysis showing regions containing tumor satellites, tumor core, NOS2^+^ edge, lymphoid aggregates, and NOS2^-^ edge.

Further examination of tumor cells localized in stroma regions provided structural details implicating progression to metastatic disease, where small clusters of elongated EpCAM positive cells that could represent metastatic phenotypes identified along NOS2^+^ edges (Figs. 5A and 6A white arrows) (13). As seen above, rich EpCAM areas show small clusters of elongated EpCAM^+^ cells in the stroma near lymphoid aggregates (Figure 5A white arrowhead) indicating tumor cell migration in the vicinity of restricted lymphoid patches that induce tumor NOS2/COX2 expression (13). In contrast to elongated EpCAM^+^ phenotypes observed near restricted lymphoid aggregates, Fig. 5E, shows CD44v6 that is expressed predominantly in the immune desert epithelial tumor core. Taken together, these results suggest that tumor NOS2/COX2 and CD8 expressions demonstrate elongated tumor NOS2/COX2 clusters that are spatially localized in stroma restricted lymphoid aggregates near the NOS2^+^ tumor edge (Fig. 7) and that tumor NOS2/COX2 expression collaborates during the metastatic process, which may involve at least in part the upregulation of EpCAM expression (26, 27).

**Figure 7.**
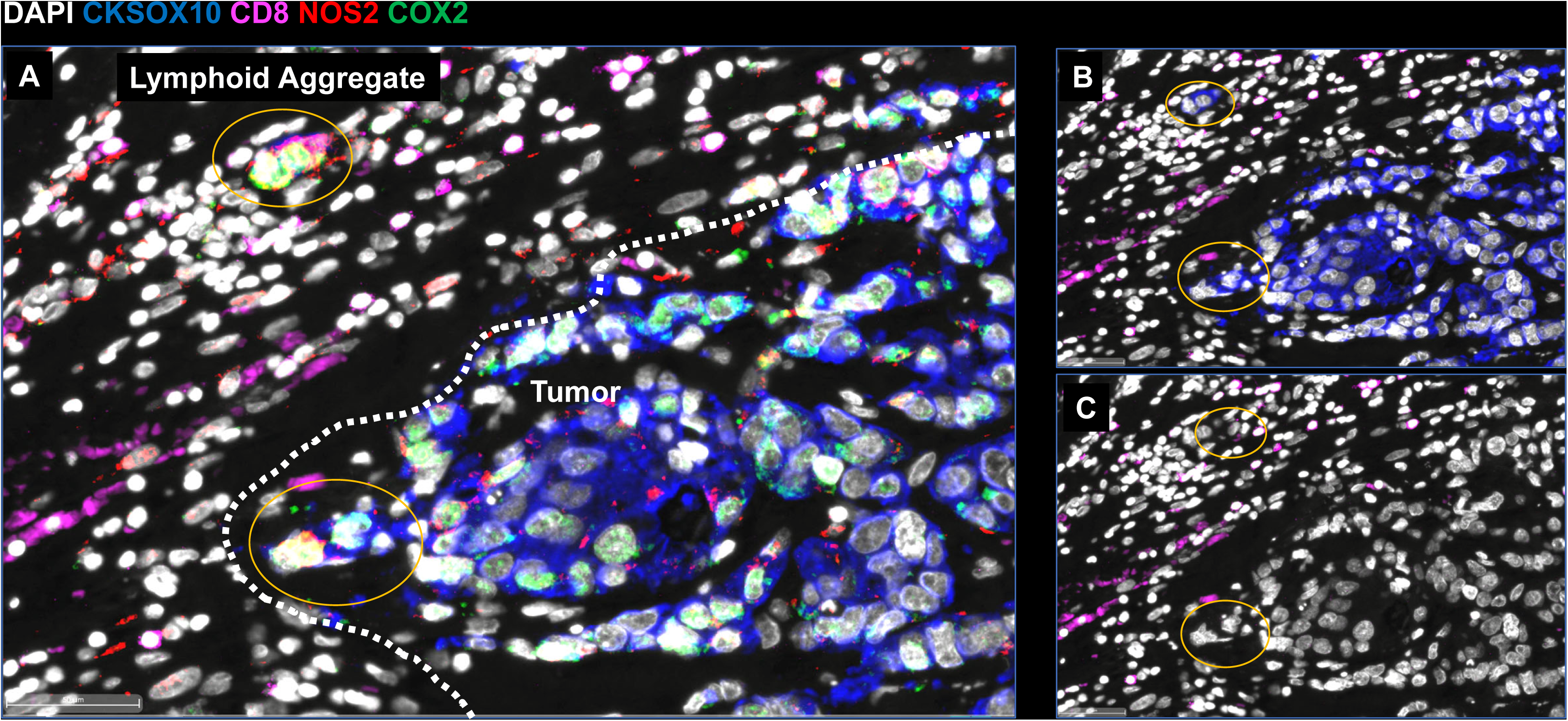
Metastatic niche showing invasive tumor edge proximal to lymphoid aggregates. A) Shows the composition of tumor marker CK-SOX10 (blue), NOS2 (red), COX2 (green) and CD8 (magenta). B) Shows the CK-SOX10 tumor marker alone relative to CD8^+^ T cell aggregate. C) DAPI with CD8 expression.

## Discussion

The strong association between tumor NOS2/COX2 co-expression and survival in ER-breast cancer suggests that these enzymes and their products are key drivers of poor clinical outcomes (2). Previous work has shown that NOS2-derived NO and COX2-derived PGE2 collaborate in a feedforward manner to activate multiple oncogenic pathways that promote metastasis, cancer stemness, and immunosuppression, which are all markers of disease progression (2, 28). These earlier observations are extended herein by the identification of unique EpCAM^+^/CD44v6^+^ cellular neighborhoods that border NOS2/COX2 high regions. Given that EpCAM and CD44v6 are markers of metastasis and cancer stemness, these observations support the role(s) of tumor NOS2/COX2 expression key targets of disease progression in breast and other tumors (27, 29). Moreover, the orthogonal expression of tumor NOS2/COX2 in distinct cells and regions within the TME suggest that the spatial configuration of these enzymes has important roles during intercellular communication. Herein we show spatial associations between regional clustering of tumor NOS2 and specific EpCAM^+^ cellular neighborhoods along the tumor margin and in the stroma in deceased patient tumors. Elevated tumor NOS2 expression and the intracellular NO levels that activate major oncogenic pathways through S-nitrosation and non heme iron is consistent with 300 μM levels determined *in vitro* (2, 3, 23). The levels of NO critical for activation of specific pathways is dependent on three factors including the rates of NO production and consumption, as well as the frequency or clustering of NOS2 expressing cells. Previous studies have shown that the higher frequency or number of clustered NOS2 expressing cells is directly proportional to the local NO concentration (10, 22). Importantly, these regions of elevated tumor NOS2 expression is spatially consistent with observed metastatic niches defined by elevated EpCAM expression.

The regulation of tumor NOS2 in human cancer cells has been somewhat of a mystery. In murine systems, the requirement of IFNγ for tumor Nos2 expression has been shown, which was amplified by other cytokines including Il1β and TNFα (22). In comparison, human NOS2 expression was detected under the same conditions *in vitro* but was considerably lower (3-5%) when compared to murine (40-100%) Nos2 expression (13, 22). Regional NOS2 expression *in vivo* is far higher in concentration (∼30-40% supplemental). CD8^+^ T cells provide a source of IFNγ in the TME and their increased presence is generally predictive of outcome clinical outcomes (30). Herein, elevated CD8^+^ T cell/tumor ratio also correlated with improved clinical outcome. Moreover, the results herein show an important spatial correlation where the stroma restriction of CD8^+^ T and increased tumor NOS2/COX2 expression correlated with poor survival. These inflamed regions of stroma restricted CD8^+^ T cells create transient areas of locally increased NO/PGE2, which promote increased oncogenic signaling (2). While tumor infiltrating CD8^+^ T cells augment therapeutic efficacies, their spatial restriction in tumor stroma is predictive of poor survival, where increases IFNγ and cytokine stimulated tumor NOS2/COX2 expression creates a cellular configuration culminating in abated CD8+ T cell infiltration as well as the development of metastatic and cancer stem cell niches (12, 13). In contrast, tumors from surviving patients at 5-yr post diagnosis exhibited low, sporadic tumor NOS2/COX2 expression and elevated CD8^+^ cell infiltration into the tumor, which promotes tumor eradication by perforins and granzyme B in a cell-to-cell contact manner. Therefore, the relationship between elevated tumor NOS2/COX2 expression and abated CD8^+^ T cell tumor infiltration implicates the importance of spatial biology. Importantly, CD8^+^ T cell restriction provides a therapeutic barrier that induces regional tumor NOS2/COX2 expression, metastasis, and cancer stemness.

Examination of different cellular niches demonstrated unexpected regional differences of CSC markers EpCAM and CD44v6, where EpCAM^+^ cells were spatially aligned with NOS2 expressing tumor cells near stroma restricted lymphoid aggregates at the tumor margin or in the tumor stroma. In contrast, CD44v6 was expressed in immune desert regions in the tumor core surrounded by COX2. EpCAM and CD44v6 can be induced by several factors including cytokines. While EpCAM can be induced by IL8 and IFNγ is inhibited by TNFα and IL6. In contrast, IL-6 and TNFα can induce CD44v6, which could in part explain the distinct EpCAM and CD44v6 spatial localization. Previously we showed IL-8 was induced in cancer cells by IFNγ and NOS2 (4, 13). In contrast, CD44v6 in some cancer is inhibited by IFNγ but induced by TNFα, IL1α, IL1β, and IL6 (31). It was also shown that IL1, TNFα induced by NO can in separate cells induce COX2 (2, 13). Also, COX2-derived PGE2 induces IL-6 (2). Taken together the differential response to cytokines associated with tumor NOS2/COX2 expression can in part explain the spatially distinct expression of CSC markers: EpCAM was increased in inflamed NOS2^+^ edge while COX2^+^ regions surrounded CD44v6 expressing cells in immune desert regions.

In summary, these results demonstrate spatially distinct tumor NOS2/COX2 and CSC biomarker expression suggesting that the production of different diffusible cytokines can shape the cellular neighborhoods that drive disease progression. The above findings show that the NOS2/COX2 spatial configuration proximal to CD8^+^ T cells create cellular niches that lead to metastasis and cancer stem cell niches. The clustering of inflamed NOS2^+^/EpCAM^+^ cellular neighborhoods promotes increased regional NO flux that drives metastatic phenotypes and poor clinical outcomes. In contrast, COX2^+^/CD44v6^+^ cellular neighborhoods localized in immune desert regions of the tumor core promote immune suppression and chemoresistant tumor phenotypes. Importantly, the characterization of these phenotypes not only have strong predictive power but can be used to design novel therapies for improved clinical outcomes.

## Materials and Methods

### Tissue Collection and Immunohistochemical Analysis of Patient Tumor Sections

Tumor specimens (n = 21) were obtained from breast cancer patients recruited at the University of Maryland (UMD) Medical Center, the Baltimore Veterans Affairs Medical Center, Union Memorial Hospital, Mercy Medical Center, and the Sinai Hospital in Baltimore between 1993 and 2003. Informed consent was obtained from all patients. The collection of tumor specimens, survey data, and clinical and pathological information (UMD protocol no. 0298229) was reviewed and approved by the UMD Institutional Review Board (IRB) for the participating institutions. The research was also reviewed and approved by the NIH Office of Human Subjects Research (OHSR no. 2248). Breast tumor NOS2 and COX2 expression was analyzed previously by IHC using 1:250 diluted NOS2 antibody and 1:50 diluted COX2 antibody (no. 610328 and 610204, respectively, BD Biosciences, San Diego, CA,) and scored by a pathologist (4, 14). For NOS2 staining, a combination score of intensity and distribution were used to categorize the immunohistochemical NOS2 stains where intensity received a score of 0-3 if the staining was negative, weak, moderate, or strong. The NOS2 distribution received scores of 0-4 for distributions <10%, 10-30%, >30-50%, >50-80% and >80% positive cells (4). For COX2 staining, scores of negative to weak (1–2) or moderate to strong (3–4) were categorized as low or high, respectively (14). Herein, NOS2 and COX2 expressions were also analyzed by fluorescent staining performed on the Leica Biosystems (Wetzlar, Germany) Bond RX Autostainer XL ST5010 using the Bond Polymer Refine Kit (Leica Biosystems DS9800), with omission of the Post Primary Block reagent, DAB and Hematoxylin. After antigen retrieval with EDTA (Bond Epitope Retrieval 2), sections were incubated for 30 min with COX2 (Cell Signaling Technology, Danvers, MA, no. 12282, 1:100), followed by the Polymer reagent and OPAL Fluorophore 520 (AKOYA, Marlborough, MA). The COX2 antibody complex was stripped by heating with Bond Epitope Retrieval 2. Sections were then incubated for 30 min with NOS2 antibody (Abcam no. ab15323, 1:50), followed by the Polymer reagent and OPAL Fluorophore 690. The NOS2 antibody complex was stripped by heating with Bond Epitope Retrieval 2 and then stained with CD8 (Abcam no. 101500, 1:100) or IFNγ (Abcam no. 231036, 1:200), followed by the Polymer reagent and OPAL Fluorophore 570. Sections were stained with DAPI and coverslipped with Prolong Gold AntiFade Reagent (Invitrogen). Images were captured using the Aperio ScanScope FL whole slide scanner (Leica). The original IHC previously reported (4, 14) and fluorescent NOS2/COX2 staining results were generally consistent.

Formalin-fixed paraffin embedded (FFPE) tissue sectioned at 4 μm and mounted on SuperFrost Plus slides were stained with a FixVUE Immuno-8^TM^ Kit (formerly referred to as UltiMapper® kits (Ultivue Inc., Cambridge, MA), USA; CD8, NOS2, COX2, CKSOX10, and IFNγ cocktail) using the antibody conjugated DNA-barcoded multiplexed immunofluorescence (mIF) method (1). These kits include the required buffers and reagents to run the assays: antibody diluent, pre-amplification mix, amplification enzyme and buffer, fluorescent probes and corresponding buffer, and nuclear counterstain reagent. Hematoxylin and Eosin (H&E) and mIF staining was performed using the Leica Biosystems BOND RX Autostainer. Before performing the mIF staining, FFPE tissue sections were baked vertically at 60-65 °C for 30 min to remove excess paraffin prior to loading on the BOND RX. The BOND RX was used to stain the slides with the recommended FixVUE (UltiMapper) protocol. During assay setup, the reagents from the kit were prepared and loaded onto the Autostainer in Leica Titration containers. Solutions for epitope retrieval (ER2, Leica Biosystems cat# AR9640), BOND Wash (Leica Biosystems cat# AR9590), along with all other BOND RX bulk reagents were purchased from Leica). During this assay, the sample was first incubated with a mixture of all 4 antibody conjugates, next the DNA barcodes of each target were simultaneously amplified to improve the sensitivity of the assay. Fluorescent probes conjugated with complementary DNA barcodes were then added to the sample to bind and label the targets; Next, a gentle signal removal step was used to remove the fluorescent probes of the markers. The stained slides were mounted in Prolong Gold Anti-Fade mountant (Thermo Fisher Scientific, Waltham, MA, cat# P36965 and coverslipped (Fisherbrand Cover Glass 22 x 40mm, #1.5). Digital immunofluorescence images were scanned at 20× magnification. Images were co-registered and stacked with Ultivue UltiStacker software. The digital images were then analyzed using HALO image analysis platform (32).

### Genome Expression Omnibus

The GSE37751 breast cancer data was obtained from the Genome Expression Omnibus (GEO) public data repository (https://www.ncbi.nlm.nih.gov/geo/info/download.html). The R software (version 4.2) was used to extract gene expression data from ER-samples for subsequent analysis. High (red) and low (black) NOS2/CD8A and COX2/CD8A ratios dichotomized at the median were calculated. The final survival data were exported to PRISM (version 9) for the purpose of creating survival graphs and statistics.

### *In vivo* studies

Animal care was provided at the NCI-Frederick Animal Facility according to procedures outlined in the Guide for Care and Use of Laboratory Animals. Our facility is accredited by the Association for Accreditation of Laboratory Animal Care International and follows the Public Health Service Policy for the Care and Use of Laboratory Animals. Female BALB/c mice obtained from the Frederick Cancer Research and Development Center Animal Production Area were used for the *in vivo* studies and housed five per cage. Eight to ten-week-old female WT and Nos2^-^ BALB/c mice were shaved a day prior to tumor injection and then were injected subcutaneously into the fourth mammary fat pad with 200,000 4T1 TNBC cells. Tumor measurements began one week after tumor cell injection, using a Vernier caliper and calculated in cubic millimeter volumes according to the following equation

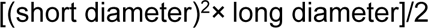

Upon reaching tumor size of 100mm^3^, tumor-bearing mice were divided into groups and treatment with 30 mg/L indomethacin in drinking water was initiated. The water was changed every Monday Wednesday Friday and treatment continued for the duration of the experiment unless otherwise specified. Experimental end point varied according to the study. Mice were euthanized at or before the tumors reach 2000mm^3^ size.

### Spatial UMAP and Neighborhood Analysis

The spatial UMAP and neighborhood analysis used here extends the analysis previously described by Giraldo, et al (33). Using NOS2 moderate and strong signal intensities, COX2 moderate signal intensities, and CD8 phenotypes (eight total phenotypes), each cell was analyzed and a phenotype density census, or neighborhood profile, was calculated using all nearby cells within specific distance ranges of 0-25 µm, 25-50 µm, 50-100µm, 100-150 µm, 150-200 µm. The neighborhood profiles of all cells in all samples (N = 1,263,845 cells total) were dimensionally reduced by a Uniform Manifold Approximation and Projection (UMAP). Clusters of similar neighborhoods were identified in the 2D representation by adaptive k-means clustering which identified 17 clusters in the UMAP. Average neighborhoods were calculated based on selected cells (e.g., cells belonging to a particular cluster). Comparisons between cluster populations were based on 5 yr outcome. Analyses were performed using MATLAB custom software and the Uniform Manifold Approximation and Projection (UMAP) library (Meehan et al https://www.mathworks.com/matlabcentral/fileexchange/71902), MATLAB Central File Exchange). Data are plotted as mean ± SEM.

### Statistical Analysis

Experiments were assayed in triplicate unless otherwise stated. Student t test was test was employed to assess statistical significance using the GraphPad Prism software (version 9). Image analyses are reported as mean ± SEM and T tests with Welch’s or Mann Whitney correction were used when appropriate to determine significance. Linear analyses and Pearson’s correlations were also conducted to determine significant correlations between protein expressions using Prism software. Significance is reported as *p ≤ 0.05, **p ≤ 0.01, ***p ≤ 0.001, ****p≤0.0001. Single cell correlation analyses were conducted in RStudio using the corrplot (0.92) in R (4.2.1).

## Acknowledgements

This project was funded in whole or in part with Federal funds from the Intramural Research Program of the NIH, National Cancer Institute, CCR, CIL (LAR, RYSC, NK, ELF, ALG, LC, RJK, SL, SMH, DWM, SKA, SA, DAW). This project has been funded in part with Federal funds from the Frederick National Laboratory for Cancer Research, National Institutes of Health, under contract HHSN261200800001E (WFH, ALW, MP, FI, DB, EFE, SKA, SJL) and Basic Science Program, Frederick National Laboratory for Cancer Research, Frederick, MD 21702 (SKA). This project was funded in part by São Paulo Research Foundation (FAPESP) grants 2018/08107-2 and 2021/14642-0 (LC, MCR), NIH R01CA238727, NIH U01CA253553, and John S Dunn Research Foundation (STCW), NCI grant no. U54 CA210181, the Breast Cancer Research Foundation (BCRF), the Moran Foundation, Causes for a Cure, philanthropic support from M. Neal and R. Neal, and the Center for Drug Repositioning and Development Program (CREDO) (JCC), Science Foundation Ireland (SFI) grant number 17/CDA/4638, and a SFI and European Regional Development Fund (ERDF) grant number 13/RC/2073 (SAG). We wish to that São Paulo Research Foundation (FAPESP) for sending students abroad (LC). The content of this publication does not necessarily reflect the views or policies of the Department of Health and Human Services, nor does mention of trade names, commercial products, or organizations imply endorsement by the US Government.

## Data Availability

Single cell RNAseq data will be made available upon request.

## Supplement figure legends

**Supplement Figure 1. Tumor analysis of NOS2 and COX2.**

A-B) Tumor NOS2 and COX2 expression at thresholds discerning strong, moderate, and weak signal intensities in Deceased and Alive patient tumors. C) Average cell intensity for NOS2 and COX2 was compared in Deceased and Alive patient tumors. * Mann Whitney p < 0.05.

**Supplement Figure 2.**

A) Upper panel showing quantification of immune and tumor (CK-SOX10) markers expressed in Deceased and Alive patient tumors. Lower panel INDO treatment upregulates mRNA expression of the thranscription factor IRF8, CLEC9a involved in anti-tumor immunity, chemokines CXCL9-11 that promote directional migration of immune cells, and IL27, which synergizes with IL12 to promote IFNγ production by CD4^+^, CD8^+^ T cells, and NKT cells. B) Comparison of whole tumor phenotypes in Deceased vs Alive patient tumors. C) Distribution plots of NOS2 vs CD8^+^ T cells of all samples showing distinct clustering of regional immune deserts, fully inflamed penetrating CD8^+^ T cells, and inflamed stroma restricted CD8^+^ T cells. D) Shows linear relationship between NOS2s and CD8 expression.

**Supplemental Figure 3. CD8^+^ T cell infiltration analysis in CD8^+/-^NOS2^+/-^COX2^+/-^ phenotypes.**

A) Deceased tumor showing inflamed stromal restricted CD8^+^ T cells with limited tumor penetration at tumor-stroma interface in a whole tumor phenotype CD8^+^NOS2^+^COX2^+^. B) Deceased tumor showing an immune desert lacking CD8^+^ T cells in a whole tumor phenotype of CD8^-^NOS2^+^COX2^+^. C) Alive tumor showing fully inflamed CD8^+^ T cell penetration in phenotype of CD8^+^NOS2^-^COX2^+/-^.

**Supplement Figure 4.**

Regional distribution of phenotype of A) CD3CD8, B) NOS2 and COX2 in tumor and macrophage with <1% Macrophage-NOS2.

