## Supplementary figures and images for "NOS2 and COX2 Provide Key Spatial Targets that Determine Outcome in ER-Breast Cancer"

### Supplementary Figures

#### Comparison of NOS2 and COX2 Expression Signal Intensities

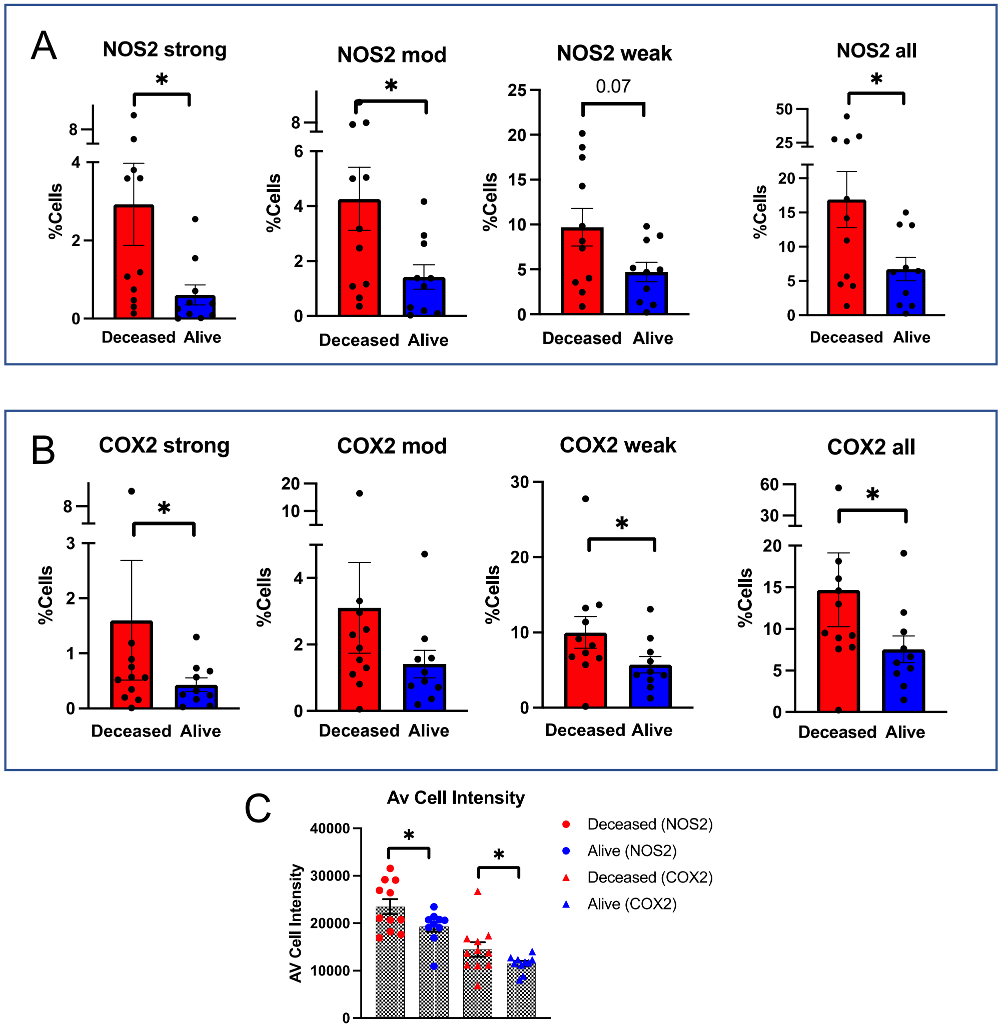


Suppl. Fig. 1

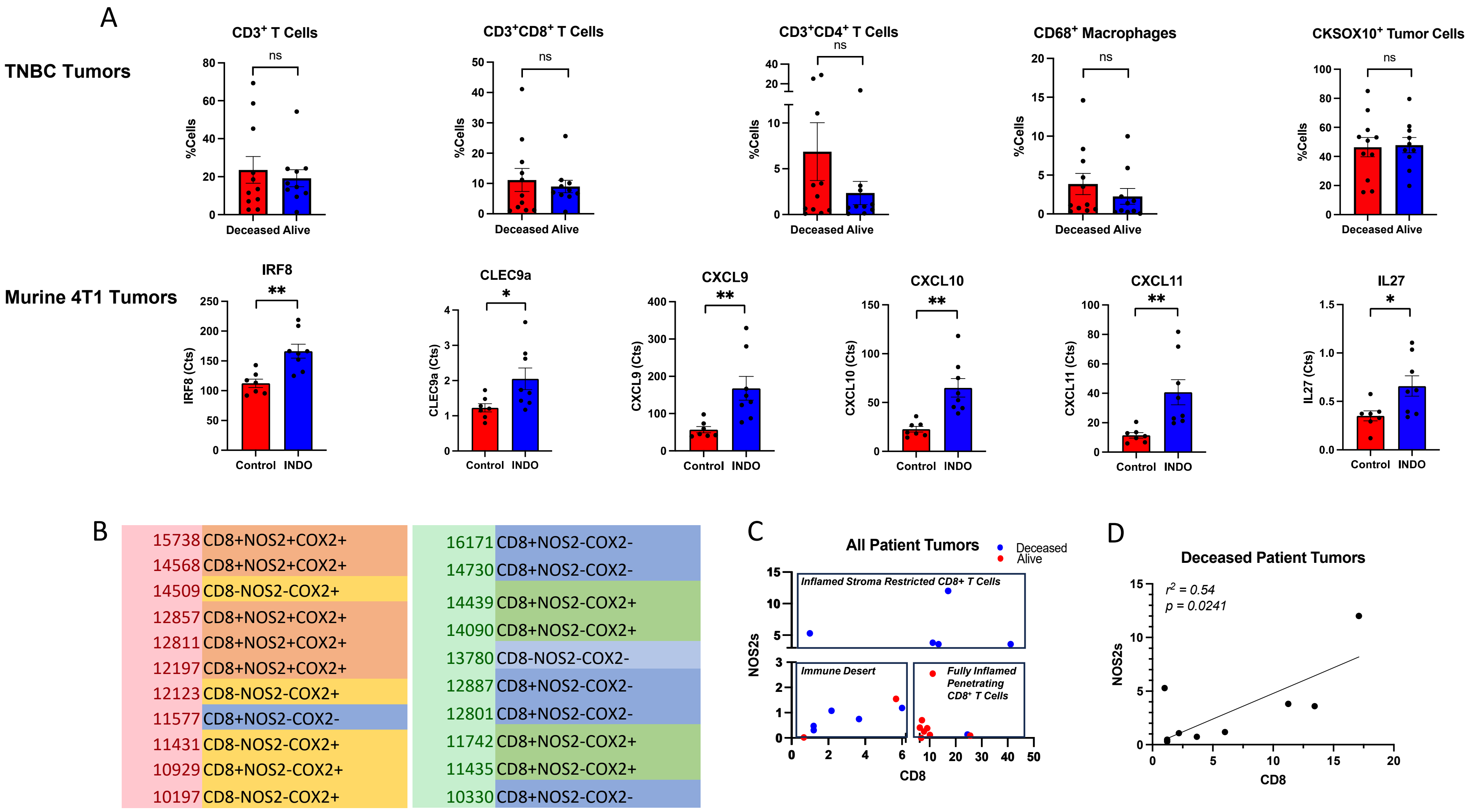

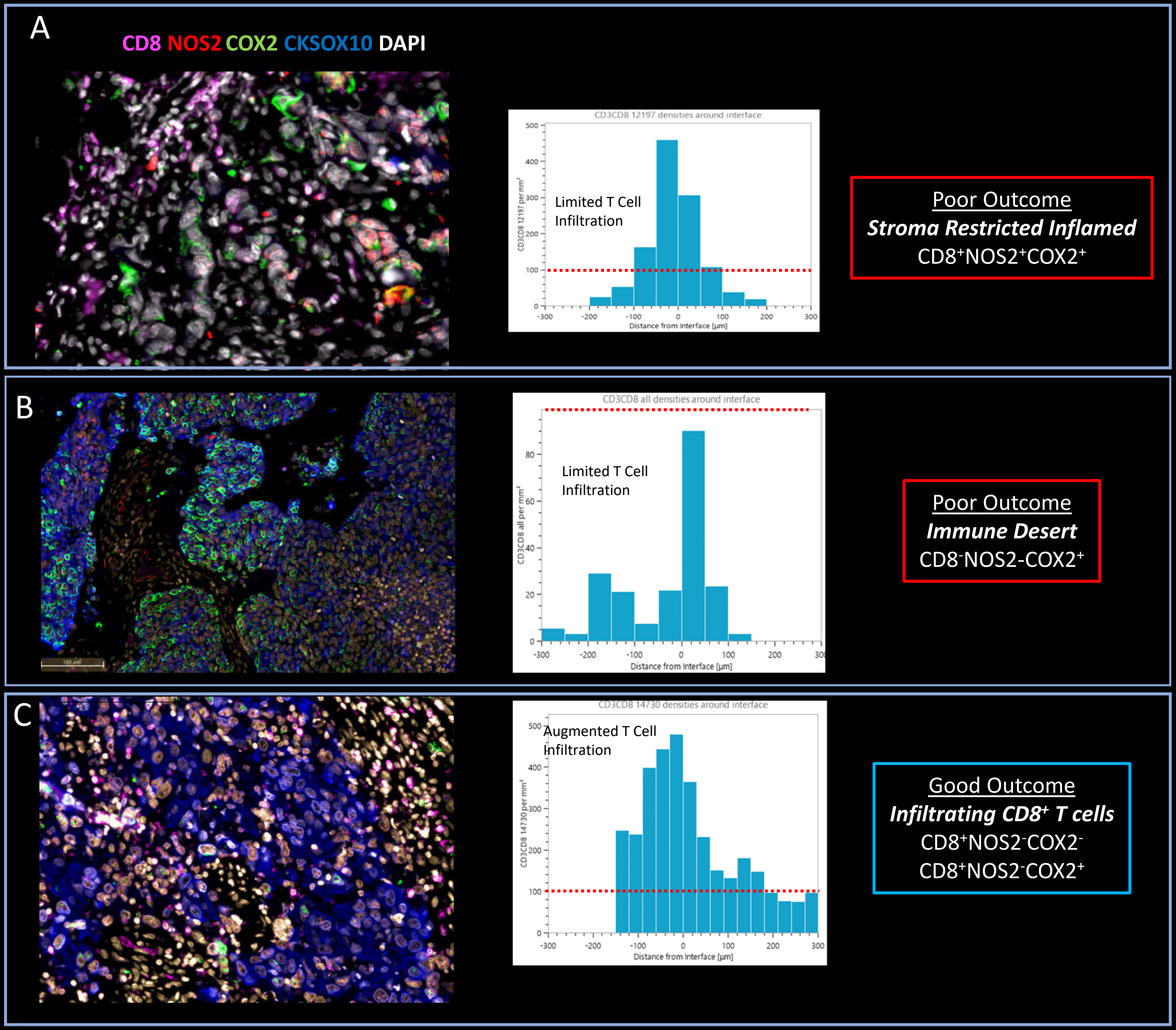

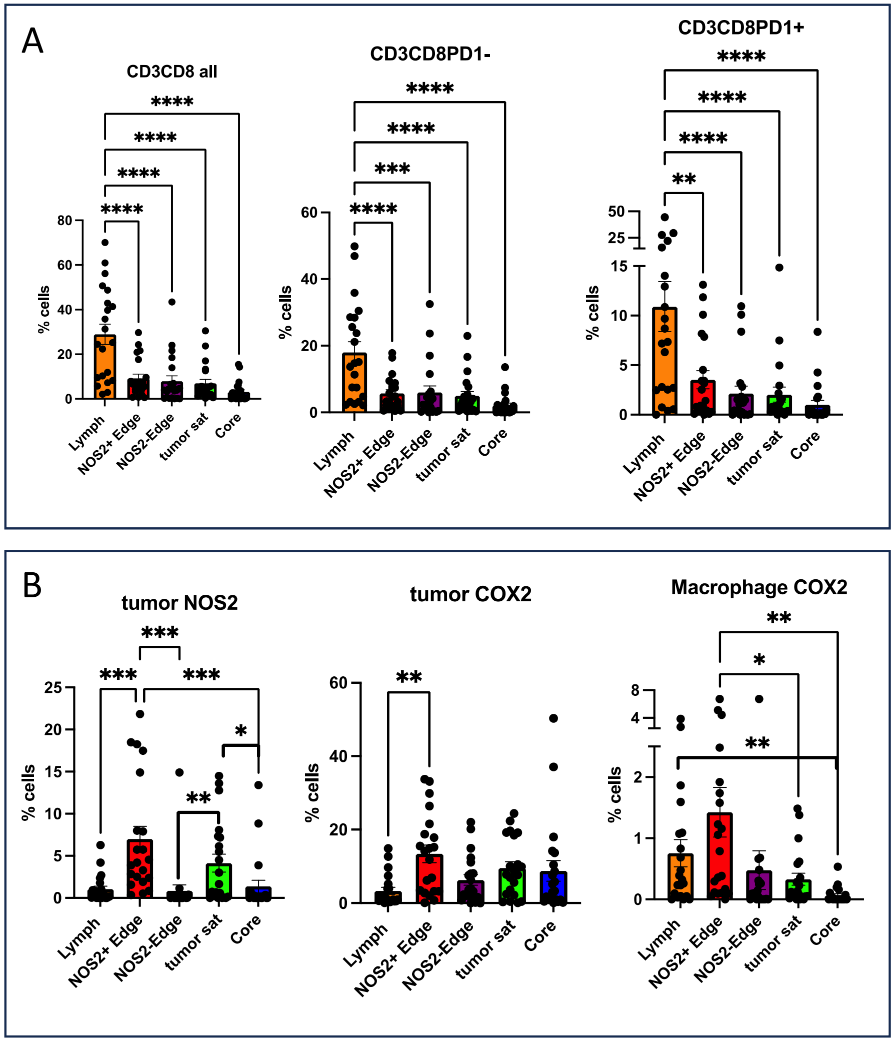
